# Genetic analysis of obstructive sleep apnoea discovers a strong association with cardiometabolic health

**DOI:** 10.1101/2020.08.04.235994

**Authors:** Satu Strausz, Sanni Ruotsalainen, Hanna M. Ollila, Juha Karjalainen, Mary Reeve, Mitja Kurki, Nina Mars, Aki S. Havulinna, Tuomo Kiiskinen, Dina Mansour Aly, Emma Ahlqvist, Maris Teder-Laving, Priit Palta, Leif Groop, Reedik Mägi, Antti Mäkitie, Veikko Salomaa, Adel Bachour, Tiinamaija Tuomi, FinnGen, Aarno Palotie, Tuula Palotie, Samuli Ripatti

**Affiliations:** Department of Oral and Maxillofacial Diseases, Helsinki University Hospital (HUH), Finland; Orthodontics, Department of Oral and Maxillofacial Diseases, Clinicum, Faculty of Medicine, University of Helsinki, Finland; Institute for Molecular Medicine Finland (FIMM/HiLIFE), University of Helsinki, Finland; Department of Psychiatry and Behavioral Sciences, Stanford University, Palo Alto, CA, USA; Analytic and Translational Genetics Unit (ATGU), Department of Medicine, Department of Neurology and Department of Psychiatry, Massachusetts General Hospital, Boston, MA, USA; Broad Institute of MIT and Harvard, Cambridge, MA, USA; Finnish Institute for Health and Welfare, Helsinki, Finland; Lund University Diabetes Centre, Department of Clinical Sciences, Malmö, Lund University, Skåne University Hospital, Malmö, Sweden; Estonian Genome Center, Institute of Genomics, University of Tartu, Tartu, Estonia; Department of Otorhinolaryngology - Head and Neck Surgery, University of Helsinki and Helsinki University Hospital, Helsinki, Finland; Research Program in Systems Oncology, University of Helsinki, Helsinki, Finland; Sleep Unit, Heart and Lung Center, Helsinki University Hospital (HUH), Finland; Endocrinology, Abdominal Centre, University of Helsinki and Helsinki University Hospital, Helsinki, Finland; Research Program for Clinical and Molecular Medicine, University of Helsinki and Folkhälsan Research Center, Helsinki, Finland; Department of Public Health, University of Helsinki, Finland

## Abstract

There is currently only limited understanding of the genetic aetiology of obstructive sleep apnoea (OSA). The aim of our study is to identify genetic loci associated with OSA risk and to test if OSA and its comorbidities share a common genetic background.

We conducted the first large-scale genome-wide association study of OSA using FinnGen Study (217,955 individuals) with 16,761 OSA patients identified using nationwide health registries.

We estimated 8.3% [0.06-0.11] heritability and identified five loci associated with OSA (P < 5.0 × 10^−8^): rs4837016 near GTPase activating protein and VPS9 domains 1 (*GAPVD1*), rs10928560 near C-X-C motif chemokine receptor 4 (*CXCR4*), rs185932673 near Calcium/calmodulin-dependent protein kinase ID (*CAMK1D*) and rs9937053 near Fat mass and obesity-associated protein (*FTO*) - a variant previously associated with body mass index (BMI). In a BMI-adjusted analysis, an association was observed for rs10507084 near Rhabdomyosarcoma 2 associated transcript (*RMST*)/NEDD1 gamma-tubulin ring complex targeting factor (*NEDD1*).

We found genetic correlations between OSA and BMI (rg=0.72 [0.62-0.83]) and with comorbidities including hypertension, type 2 diabetes (T2D), coronary heart disease (CHD), stroke, depression, hypothyroidism, asthma and inflammatory rheumatic diseases (IRD) (rg > 0.30). Polygenic risk score (PRS) for BMI showed 1.98-fold increased OSA risk between the highest and the lowest quintile and Mendelian randomization supported a causal relationship between BMI and OSA.

Our findings support the causal link between obesity and OSA and joint genetic basis between OSA and comorbidities.

## Introduction

Obstructive sleep apnoea (OSA) is a severe sleep disorder affecting at least 9% of the population. Prevalence increases with higher age reaching over 35% in individuals over 60 years of age^1^. Despite a recognized health impact and available diagnostic tools and treatments the condition remains underdiagnosed^2, 3^. OSA is characterized by repetitive episodes of nocturnal breathing cessation due to upper airway collapse resulting in mild to severe sleep deprivation and dysregulation of sleep, breathing and blood pressure. These conditions may lead to serious comorbidities through intermittent hypoxia, systemic inflammation and sympathetic activation^4^. Furthermore, OSA is influenced by multiple risk factors such as obesity, male sex, family history of OSA, high age and problems of upper airway flow or jaw anatomy^5^.

Consequently, OSA is a serious public health problem due to its many cardiometabolic comorbidities including an increased risk to coronary heart disease (CHD), type 2 diabetes (T2D) and its complications^6^ and ultimately, increased mortality^7^. In addition, comorbidities such as depression^8^, hypothyroidism^9^, asthma^10^ and inflammatory rheumatic diseases (IRD)^11^ are linked with OSA. IRD might manifest as a comorbidity of OSA through the affection of the temporomandibular joint, which rotates the lower jaw backward causing narrowing of the upper airway^12^.

Genetic studies provide a tool to identify independent genetic risk factors that modulate disease risk, and to examine causal pathways between comorbidity traits. Genome-wide association studies (GWAS) in OSA patients have previously identified associations with OSA severity measured with apnoea-hypopnea index (AHI, number of apnoeas and hypopneas per hour of sleep) or respiratory event duration^13–15^. The genome-wide significant findings from these studies and the corresponding associations our study are found in **Supplementary Table 1**. Larger-scale GWAS studies have been performed on OSA-related phenotypes such as snoring^16^. However, knowledge about OSA predisposing genetic loci is thus far limited^17^.

To test genetic associations with OSA we utilised FinnGen study with genetic profiling for 217,955 individuals and OSA diagnosis based on International Statistical Classification of Diseases (ICD) codes obtained from the Finnish National Hospital Discharge Registry and the Causes of Death Registry. The registries have excellent validity and coverage^18^. Combining the OSA diagnosis (ICD-10: G47.3, ICD-9: 3472) and related risk factors and comorbidities with the genotyping data allows identification of risk variants, helps elucidating biological disease mechanisms and enables evaluation of OSA-related disease burden on a population level.

The aim of the study is to identify genetic loci associated with OSA risk and to test if OSA and its comorbidities share a common genetic background. To our knowledge, this is the first population-level longitudinal GWAS study regarding OSA.

## Materials and Methods

### General information

First, using the FinnGen data, a GWAS was calculated for 2,925 ICD-code based phenotype definitions including OSA. Second, we selected into further analyses comorbidities which have previously been shown to associate with OSA in epidemiological studies, including obesity^19^, hypertension^20^, T2D^21^, CHD, stroke^22^, depression^8^, hypothyroidism^9^, asthma^10^ and IRD^11, 12^.

### Study sample in FinnGen

FinnGen (https://www.finngen.fi/en) is a large biobank study that aims to genotype 500,000 Finns and combine this data with longitudinal registry data including The National Hospital Discharge Registry, Causes of Death Registry and medication reimbursement registries, all these using unique national personal identification codes. FinnGen includes prospective and retrospective epidemiological and disease-based cohorts as well as hospital biobank samples. The data consists of 218,792 individuals until the spring of 2020. FinnGen’s genotyping and imputation protocol is described in **Supplementary Information**.

To examine OSA patients more specifically 837 individuals who had ICD-code G47 (Sleep disorders) were excluded from the controls and thus the remaining sample size was 217,955 participants. Of them, 16,761 (7.7%) had OSA diagnosis and 10,557 (63.0%) of OSA patients were male. Baseline characteristics and OSA comorbidities of the participants are presented in **Table 1.** Differences in baseline demographics and clinical characteristics were tested using logistic regression model. The model was adjusted for sex, age and 10 first principal components (PC), except the model for age was adjusted for sex and 10 first PCs and the model for sex was adjusted for age and 10 first PCs.

**Table 1.**
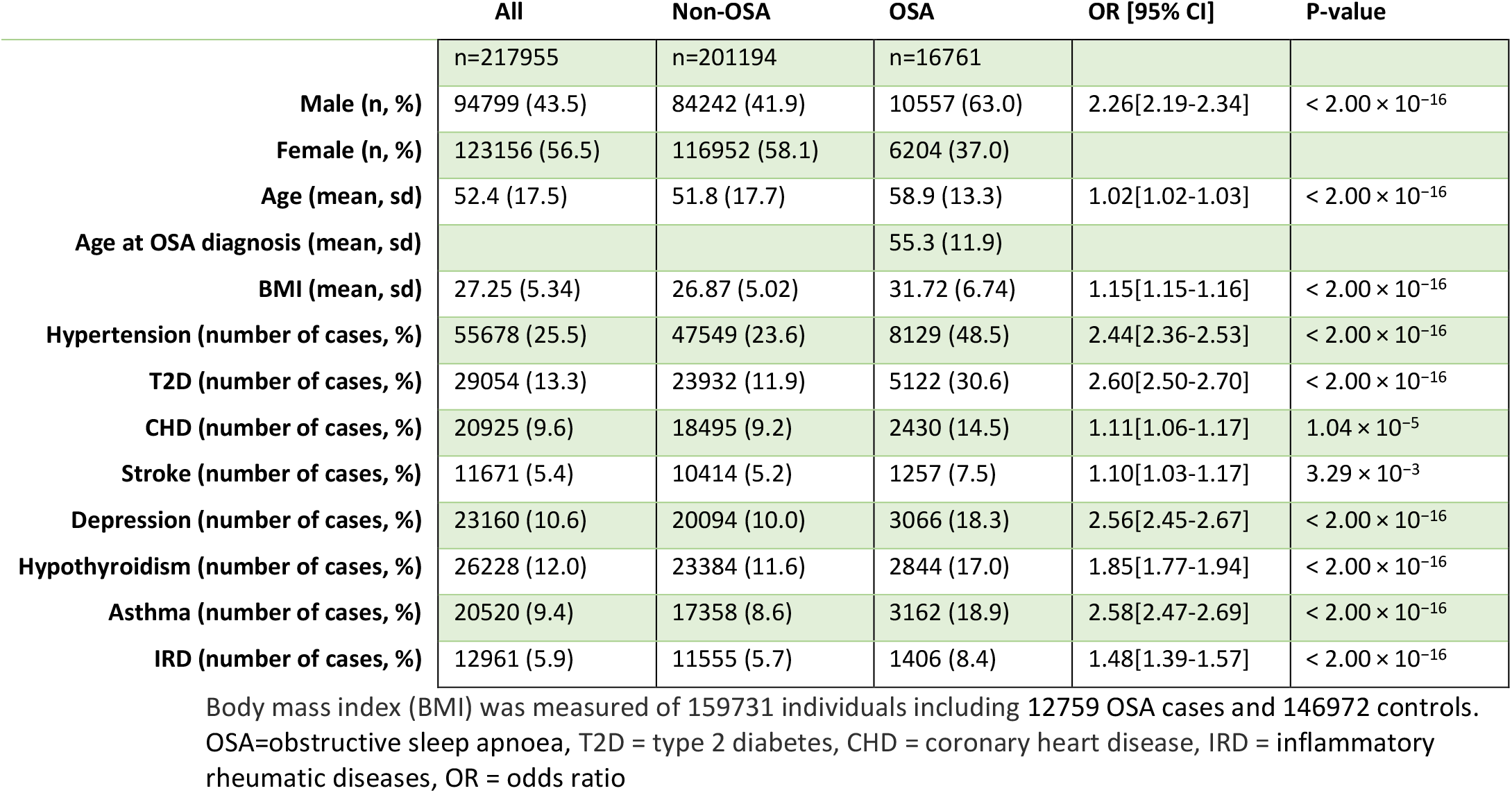
Baseline characteristics and previously known OSA comorbidities between OSA and non-OSA individuals in the FinnGen cohort

|  | All | Non-OSA | OSA | OR [95% CI] | P-value |
| --- | --- | --- | --- | --- | --- |
|  | n=217955 | n=201194 | n=16761 |  |  |
| Male (n, %) | 94799 (43.5) | 84242 (41.9) | 10557 (63.0) | 2.26[2.19-2.34] | $< 2.00 \times 10^{-16}$ |
| Female (n, %) | 123156 (56.5) | 116952 (58.1) | 6204 (37.0) |  |  |
| Age (mean, sd) | 52.4 (17.5) | 51.8 (17.7) | 58.9 (13.3) | 1.02[1.02-1.03] | $< 2.00 \times 10^{-16}$ |
| Age at OSA diagnosis (mean, sd) |  |  | 55.3 (11.9) |  |  |
| BMI (mean, sd) | 27.25 (5.34) | 26.87 (5.02) | 31.72 (6.74) | 1.15[1.15-1.16] | $< 2.00 \times 10^{-16}$ |
| Hypertension (number of cases, %) | 55678 (25.5) | 47549 (23.6) | 8129 (48.5) | 2.44[2.36-2.53] | $< 2.00 \times 10^{-16}$ |
| T2D (number of cases, %) | 29054 (13.3) | 23932 (11.9) | 5122 (30.6) | 2.60[2.50-2.70] | $< 2.00 \times 10^{-16}$ |
| CHD (number of cases, %) | 20925 (9.6) | 18495 (9.2) | 2430 (14.5) | 1.11[1.06-1.17] | $1.04 \times 10^{-5}$ |
| Stroke (number of cases, %) | 11671 (5.4) | 10414 (5.2) | 1257 (7.5) | 1.10[1.03-1.17] | $3.29 \times 10^{-3}$ |
| Depression (number of cases, %) | 23160 (10.6) | 20094 (10.0) | 3066 (18.3) | 2.56[2.45-2.67] | $< 2.00 \times 10^{-16}$ |
| Hypothyroidism (number of cases, %) | 26228 (12.0) | 23384 (11.6) | 2844 (17.0) | 1.85[1.77-1.94] | $< 2.00 \times 10^{-16}$ |
| Asthma (number of cases, %) | 20520 (9.4) | 17358 (8.6) | 3162 (18.9) | 2.58[2.47-2.69] | $< 2.00 \times 10^{-16}$ |
| IRD (number of cases, %) | 12961 (5.9) | 11555 (5.7) | 1406 (8.4) | 1.48[1.39-1.57] | $< 2.00 \times 10^{-16}$ |

Body mass index (BMI) was measured of 159731 individuals including 12759 OSA cases and 146972 controls. OSA=obstructive sleep apnoea, T2D = type 2 diabetes, CHD = coronary heart disease, IRD = inflammatory rheumatic diseases, OR = odds ratio

The diagnosis of OSA was based on ICD-codes (ICD-10: G47.3, ICD-9: 3472A), which were obtained from the Finnish National Hospital Discharge Registry and the Causes of Death Registry. This diagnosis is based on subjective symptoms, clinical examination and sleep registration applying AHI≥5/hour or respiratory disturbance index (RDI)≥5/hour. By combining ICD-codes from different registries, we generated disease endpoints. **Supplementary Table 2** describes how endpoints were constructed for each phenotype.

All prescription medicine purchases were retrieved from the Social Insurance Institution of Finland (KELA) registry for prescription drug purchases, since 1995 (excluding over-the-counter medicines and medication administered at hospitals). The drugs are coded by the Anatomical Therapeutic Chemical (ATC) Classification System.

### Study samples in other cohorts

UK Biobank (UKBB, https://www.ukbiobank.ac.uk/) is a major national and international health resource, with the aim of improving the prevention, diagnosis and treatment of a wide range of serious and life-threatening illnesses. UKBB recruited 500,000 people in 2006-2010 from across the United Kingdom. OSA diagnosis was based on ICD-10: G47.3. The study sample in the UKBB included 4,471 OSA cases and 403,723 controls.

The Estonian Biobank is a population-based biobank of the Estonian Genome Center at the University of Tartu (EGCUT, www.biobank.ee). The cohort size is currently close to 150,000 participants. Patients were selected by ICD-10: G47.3. For additional conformation of the diagnosis treatment service codes from the Health Insurance Fund were also used. The study sample in the EGCUT included 4,930 OSA patients and 61,056 controls.

The All New Diabetics in Scania (ANDIS, http://andis.ludc.med.lu.se/) aims to recruit all incident cases of diabetes within Scania County in Southern Sweden. All health care providers in the region were invited; the current registration covered 14,625 patients. OSA was defined by ICD-10: G47.3. The study sample included 947 OSA patients and 9,829 controls.

### GWAS

A total of 218,792 samples from FinnGen Data Freeze 5 with 2,925 disease endpoints were analyzed using Scalable and Accurate Implementation of Generalized mixed model (SAIGE), which uses saddle point approximation (SPA) to calibrate unbalanced case-control ratios^23^. Analyses were adjusted for age, sex, genotyping chip, genetic relationship and first 10 PCs. For OSA, we performed GWAS in a similar manner (n=217,955, including 16,761 OSA patients and 201,194 controls), but adjusting also for body mass index (BMI) (n=159,731, including 12,759 OSA patients and 146,972 controls).

For the replication of the FinnGen OSA GWAS results we merged the evidence from the UKBB, EGCUT and ANDIS cohorts. The results were combined using inverse-variance weighted fixed-effect meta-analysis. The merged data consisted 10,348 OSA cases and 474,608 controls.

The GWAS using UKBB data was calculated using SAIGE^23^. This subset included 4,471 OSA cases and 403,723 controls and was adjusted for birth year, sex, genetic relatedness and the first 4 PCs. In the EGCUT the data were analyzed using SAIGE and the model was adjusted for age, sex, genetic relatedness and the first 10 PCs and included 4,930 OSA patients and 61,056 controls. In ANDIS, the GWAS was calculated using logistic regression model, which was adjusted for age, sex and first 10 PCs. The analysis included 947 cases and 9,829 controls.

### Linkage disequilibrium score regression (LDSC)

To estimate single nucleotide polymorphism (SNP) -based heritability, genetic correlation and tissue specific SNP-heritability we used LDSC-software^24^. LDSC uses linkage disequilibrium (LD) score regression method, which quantifies the contribution of each variant by examining the relationship between test statistics and LD. In calculation we used LD scores calculated from the 1000 Genomes Project^25^. To restrict to a set of common, well-imputed variants, we retained only those SNPs in the HapMap 3 reference panel^26^.

To study genetic correlations between OSA, BMI, hypertension, T2D, CHD, stroke, depression, hypothyroidism, asthma and IRD we used summary statistics from the FinnGen data. For sleep traits we used summary statistics derived from the UKBB data. Study subjects self-reported sleep duration, sleepiness^27^ and chronotype^28^. Sleep efficiency (sleep duration divided by the time between the start and end of the first and last nocturnal inactivity period, respectively) was based on accelerometer-derived measures^29^. For tissue specific SNP-heritability we used a method, which combined data from Encyclopedia of DNA Elements (ENCODE, https://www.encodeproject.org/) and the Genotype-Tissue Expression (GTEx, https://gtexportal.org/home/) resources^30, 31^.

### Polygenic risk score (PRS) and Mendelian randomization (MR)

PRS for BMI was calculated using summary statistics for 996,250 variants^32^. The posterior effect sizes were calculated with PRS-CS utilising method^33^ and the score was calculated using Plink2 (https://www.cog-genomics.org/plink/2.0/) for the FinnGen data.

We performed MR analysis to investigate the causality between BMI and OSA using independent BMI SNPs^32^. A genetic variant associated with the exposure of interest (genetic instrument) is used to test the causal relationship with the exposure (BMI) and outcome (OSA)^34^.

### Gene based analysis

Gene-based tests were performed using Multi-marker Analysis of GenoMic Annotation (MAGMA) as implemented on the Functional Mapping and Annotation (FUMA) platform, which provides aggregate association p-values based on all variants located within a gene and its regulatory region using information from 18 biological data repositories and tools^35^. This analysis includes a gene-based test to detect significant SNPs associated with OSA using FinnGen OSA summary statistics.

## Results

### OSA correlates strongly with cardiovascular and metabolic traits

To estimate strengths of associations between OSA and comorbidities we utilised data from 217,955 individuals who have participated in the FinnGen project. 16,761 (7.7%) had OSA diagnosis and 10,557 (63%) of cases were male. The diagnoses were derived from ICD-codes in the Finnish National Hospital Discharge Registry and from the Causes of Death Registry. Baseline characteristics of the FinnGen participants and odds for OSA associated comorbidities are presented in Table 1.

### GWAS of OSA reveals BMI dependent and independent associations

We estimated the heritability for OSA in FinnGen to be 8.3% [0.06-0.11] before and 6.0% [0.04-0.08] after BMI adjustment. In a genome-wide association test, five distinct genetic loci were associated with OSA (P < 5.0 × 10^−8^), outlined in **Table 2** and **Figure 1a** and regional associations in **Supplementary Figure 1**. The lead variant in a locus on chromosome 16 was rs9937053, an intronic variant near Fat mass and obesity-associated protein (*FTO*), P = 4.3 × 10^−16^. In chromosome 12, the lead variant was rs10507084, near Rhabdomyosarcoma 2 associated transcript (*RMST*)/NEDD1 gamma-tubulin ring complex targeting factor (*NEDD1*), P = 2.8 × 10^−11^, where *RMST*, a long non-coding RNA, was the nearest gene and *NEDD1* the nearest protein coding gene. On chromosome 10, the lead variant was rs185932673, an intronic variant near Calcium/calmodulin-dependent protein kinase ID (*CAMK1D*), P = 2.4 × 10^−8^. In chromosome 9, the lead variant was rs4837016 near GTPase activating protein and VPS9 Domains 1 (*GAPVD1*), P = 1.5 × 10^−8^ and in chromosome 2, the lead variant rs10928560 was near C-X-C motif chemokine receptor 4 (*CXCR4*), P = 2.8 x 10^−8^. Four out of five of these OSA associated lead variants have also been previously associated with BMI (p<0.01)^36–38^, with the exception of rs10507084 at the *RMST/NEDD1* locus. Conditional analyses of the associated loci did not suggest any additional associations. Adjusting for BMI did not affect the association for variant rs10507084 (**Figure 1b** and Table 2), (OR_unadjusted_ = 1.11[1.08-1.15], P=2.8 × 10^−11^ vs. OR_BMI adjusted_ = 1.12[1.08-1.17], P=9.7 × 10^−10^) suggesting BMI-independent mechanisms for rs10507084 in OSA predisposition.

**Table 2.** Characterization of five genome-wide significant OSA loci

| CHR | Position | RSID | Nearest gene | Consequence | Fin.enr. | AF | AF cases | AF controls | OR | p-value | p-value BMIadj |
| --- | --- | --- | --- | --- | --- | --- | --- | --- | --- | --- | --- |
| 16 | 53765595 | rs9937053 | FTO | intron | 0.97 | 0.43 | 0.45 | 0.43 | 1.11[1.08-1.13] | $4.3 \times 10^{-16}$ | 0.04 |
| 12 | 97359374 | rs10507084 | RMST/NEDD1 | intergenic | 3.03 | 0.18 | 0.19 | 0.18 | 1.11[1.08-1.15] | $2.8 \times 10^{-11}$ | $9.7 \times 10^{-10}$ |
| 10 | 12656440 | rs185932673 | CAMK1D | intron | 0.55 | 0.0033 | 0.0051 | 0.0032 | 1.87[1.50-2.33] | $2.4 \times 10^{-8}$ | $9.3 \times 10^{-6}$ |
| 9 | 125379530 | rs4837016 | GAPVD1 | intergenic | 1.12 | 0.47 | 0.45 | 0.47 | 0.93[0.91-0.95] | $1.5 \times 10^{-8}$ | $2.2 \times 10^{-4}$ |
| 2 | 136234237 | rs10928560 | CXCR4 | downstream | 1.04 | 0.20 | 0.18 | 0.20 | 0.92[0.89-0.94] | $2.8 \times 10^{-8}$ | $8.5 \times 10^{-5}$ |

The finding of rs185932673 should be interpreted cautiously as the variant is rare in the Finnish population. CHR=chromosome, Fin.enr=Finnish enrichment compared to other European ancestry, AF=allele frequency, OR=odds ratio, [95% confidence interval], p-value BMIadj=p-value after BMI adjustment

**Figure 1.**
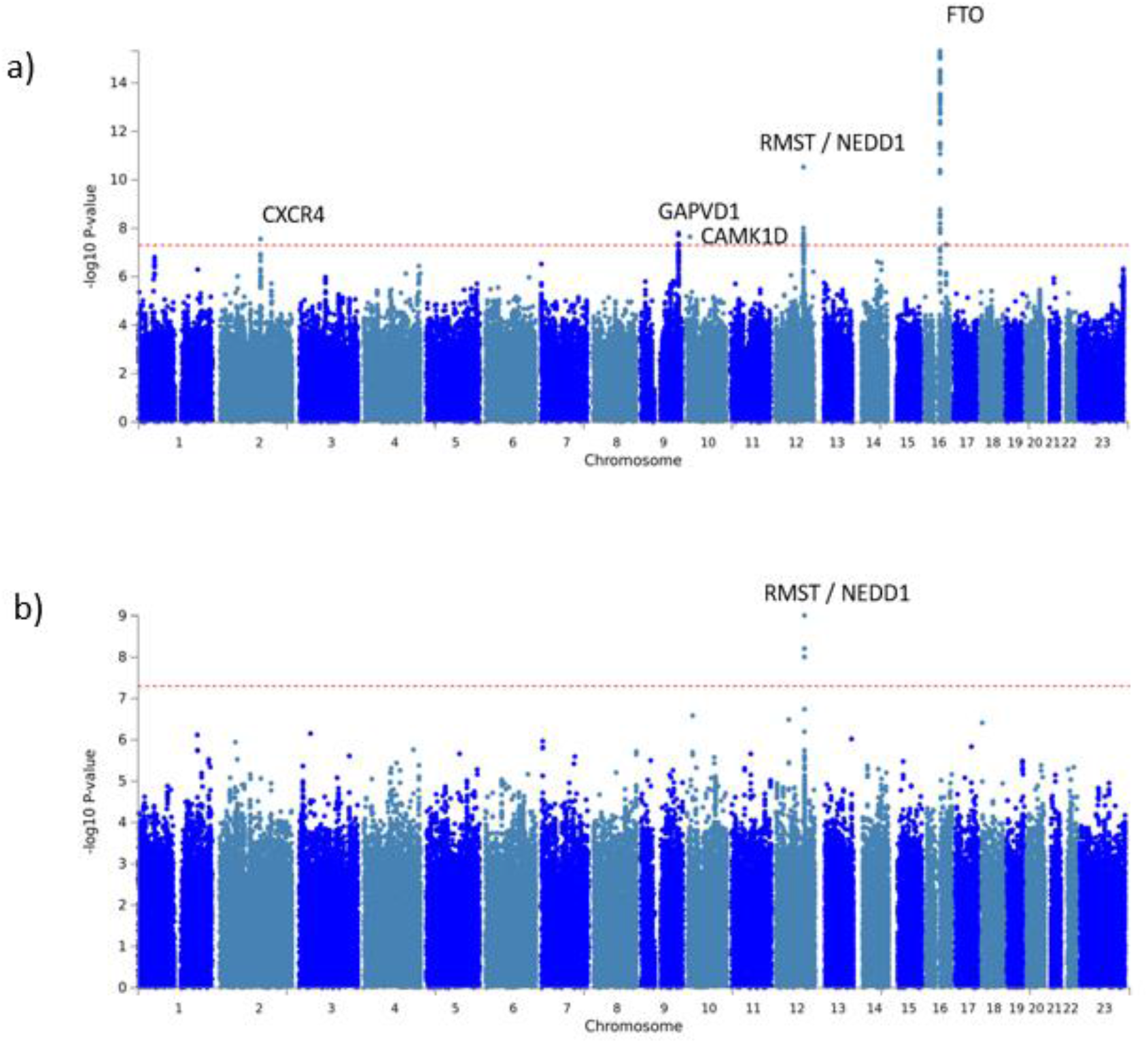
a) Manhattan plot for obstructive sleep apnoea (OSA) including 16 761 OSA cases and 201 194 controls. For each genetic variant, the x-axis shows chromosomal position, while y-axis shows the −log_10_(P) value. The horizontal line indicates the genome-wide significance threshold of P = 5 × 10^−8^. Five genetic loci were identified at the genome-wide significance level. *CXCR4=C-X-C* motif chemokine receptor 4, *GAPVD1=* GTPase activating protein and VPS9 Domains 1, *CAMK1D=*Calcium/calmodulin-dependent protein kinase ID, *RMST*=Rhabdomyosarcoma 2 associated transcript / *NEDD1*=NEDD1 gamma-tubulin ring complex targeting factor, *FTO=Fat* mass and obesity-associated protein b) Manhattan plot for obstructive sleep apnoea (OSA) after body mass index (BMI) adjustment including 12 759 OSA cases and 146 972 controls. For each genetic variant, the x-axis shows chromosomal position, while y-axis shows the −log_10_(P) value. The horizontal line indicates the genome-wide significance threshold of P = 5 × 10^−8^. One genetic locus was identified at the genome-wide significance level. *RMST*=Rhabdomyosarcoma 2 associated transcript / *NEDD1*=NEDD1 gamma-tubulin ring complex targeting factor

As an exploratory analysis we used MAGMA. This tool annotates FinnGen OSA summary statistics based on 18 biological data repositories and tools^35^. Using MAGMA, we detected 25 significant associations (P < 2.54 × 10^−6^) with various biological processes, which were driven by the same loci as the significant GWAS variants in *FTO* and *GAPVD1* **(Supplementary Figure 2a)**. Similarly, the gene-based test for BMI-adjusted OSA revealed three further associated genes (**Supplementary Figure 2b)**.

We performed a phenome-wide association analysis (PheWAS) using the FinnGen data and examined the associations between the lead SNPs and 2,925 disease endpoints. Rs10507084 was specific for OSA also after BMI adjustment suggesting an independent role from cardiometabolic traits for the association between rs10507084 and OSA (**Figure 2a**). In addition, there was a strong correlation between rs10507084 and the use of antidepressants (OR=1.013[1.007-1.019], P=4.4 × 10^−6^) (**Figure 2b**).

**Figure 2.**
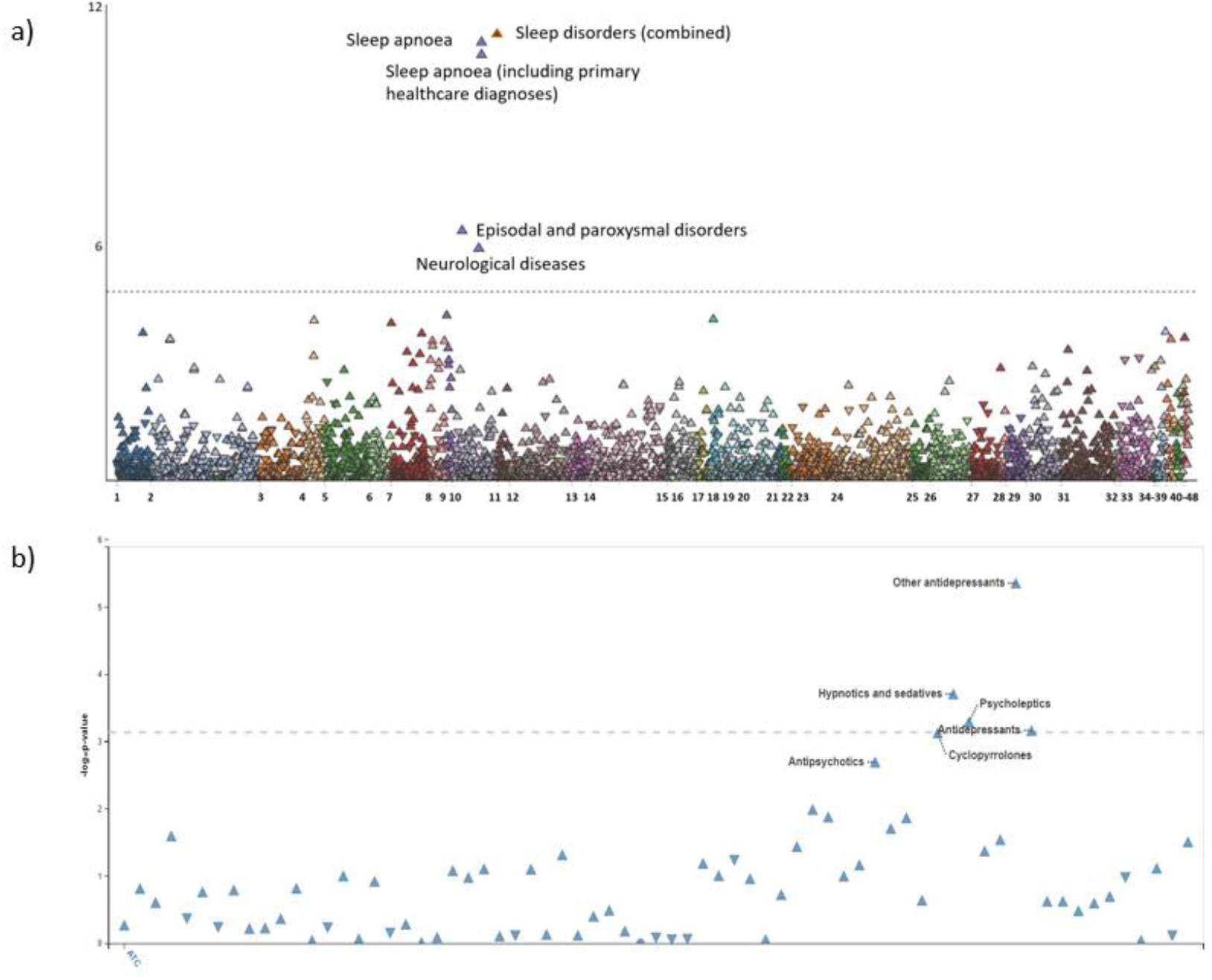
a) Phenome-wide association analysis (PheWAS) associations after body mass index (BMI) adjustment between rs10507084 and 2,925 disease endpoints. Significance Bonferroni corrected threshold was defined at P = 0.05/2925 = 1.71 × 10^−5^. Associated P-values on the −log_10_ scale on the vertical axis and the disease definition along the horizontal axis: 1. I Certain infectious and parasitic diseases, 2. II Neoplasms from hospital discharges, 3. II Neoplasms, from cancer registry, 4. III Diseases of the blood and blood-forming organs and certain disorders involving the immune mechanism, 5. IV Endocrine, nutritional and metabolic diseases, 6. Diabetes endpoints, 7. V Mental and behavioural disorders, 8. Psychiatric endpoints, 9. Alcohol related diseases, 10. VI Diseases of the nervous system, 11. Neurological endpoints, 12. VII Diseases of the eye and adnexa, 13. VIII Diseases of the ear and mastoid process, 14. IX Diseases of the circulatory system, 15. Cardiometabolic endpoints, 16. X Diseases of the respiratory system, 17. Asthma and related endpoints, 18. Chronic obstructive pulmonary disease and related endpoints, 19. Interstitial lung disease endpoints, 20. XI Diseases of the digestive system, 21. Dental endpoints, 22. Gastrointestinal endpoints, 23. XII Diseases of the skin and subcutaneous tissue, 24. XIII Diseases of the musculoskeletal system and connective tissue, 25. Rheumatoid arthritis endpoints, 26. XIV Diseases of the genitourinary system, 27. XV Pregnancy, childbirth and the puerperium, 28. XVI Certain conditions originating in the perinatal period, 29. XVII Congenital malformations, deformations and chromosomal abnormalities, 30. XVIII Symptoms, signs and abnormal clinical and laboratory findings, not elsewhere classified, 31. XIX Injury, poisoning and certain other consequences of external causes, 32. XX External causes of morbidity and mortality, 33. XXI Factors influencing health status and contact with health services, 34. Drug purchase endpoints, 35. Diseases marked as autoimmune origin, 36. Common endpoint, 37. Demonstration endpoints, 38. ICD-10 main chapters, 39. Operation endpoints, 40. Other, not yet classified endpoints, 41. Miscellaneous, not yet classified endpoints, 42. Comorbidities of Asthma, 43. Comorbidities of Chronic obstructive pulmonary disease, 44. Comorbidities of Diabetes, 45. Comorbidities of Gastrointestinal endpoints, 46. Comorbidities of Interstitial lung disease endpoints, 47. Comorbidities of Neurological endpoints, 48. Comorbidities of Rheumatoid arthritis endpoints b) Phenome-wide association analysis (PheWAS) analysis concerning drug purchases. The x-axis shows phenotypes based on Anatomical Therapeutic Chemical – drug codes (ATC), while y-axis shows the significance Bonferroni corrected threshold −log_10_(P) value which was defined as 0.05/69 = 7.25 × 10^−4^.

### Genetic correlations and MR connect OSA with cardiovascular outcomes and dysregulation of metabolism

To study the potential common genetic background of OSA and its known epidemiological correlates, we computed genetic correlations between OSA and its comorbidities using FinnGen summary statistics. The results showed strong genetic correlations between OSA and BMI (rg = 0.72, [0.62-0.83], P=3.49 × 10^−40^) and between OSA and comorbidities: hypertension (rg=0.35, [0.23-0.48], P=4.06 × 10^−8^), T2D (rg=0.52, [0.37-0.66], P=6.40 × 10^−12^), CHD (rg=0.38, [0.17-0.58], P=3.84 × 10^−4^), stroke (rg=0.33, [0.03-0.63], P=2.93 × 10^−2^), depression (rg=0.43, [0.27-0.60], P=2.79 × 10^−7^), hypothyroidism (rg=0.40, [0.27-0.54], P=7.07 × 10^−9^), asthma (rg=0.50, [0.33-0.68], P=1.53 × 10^−8^) and IRD (rg=0.34, [0.09-0.58], P=6.97 × 10^−3^). Furthermore, we observed high genetic correlations between OSA comorbidities. Since many of OSA comorbidities are correlated with BMI, we calculated the genetic correlations after BMI adjustment. This analysis showed somewhat lower estimates for genetic correlations between OSA and CHD (rg=0.24 [0.012-0.47], P=0.04), depression (rg=0.33, [0.17-0.50], P=1.1 × 10^−3^), asthma (rg=0.33 [0.11-0.54], P=2.6 × 10^−3^) and hypothyroidism, (rg=0.28 [0.11-0.44], P=8.0 × 10^−4^). Genetic correlations between OSA and BMI (rg=0.08, [−0.05-0.22], P=0.22), hypertension (rg=0.05, [−0.10-0.20], P=0.51), T2D (rg=0.15, [−0.03-0.33], P=0.11), stroke (rg=0.32, [−0.05-0.69], P=0.09) and IRD (rg=0.27, [−0.01-0.54], P=5.7 × 10^−2^) attenuated after BMI adjustment (**Figure 3)**.

**Figure 3.**
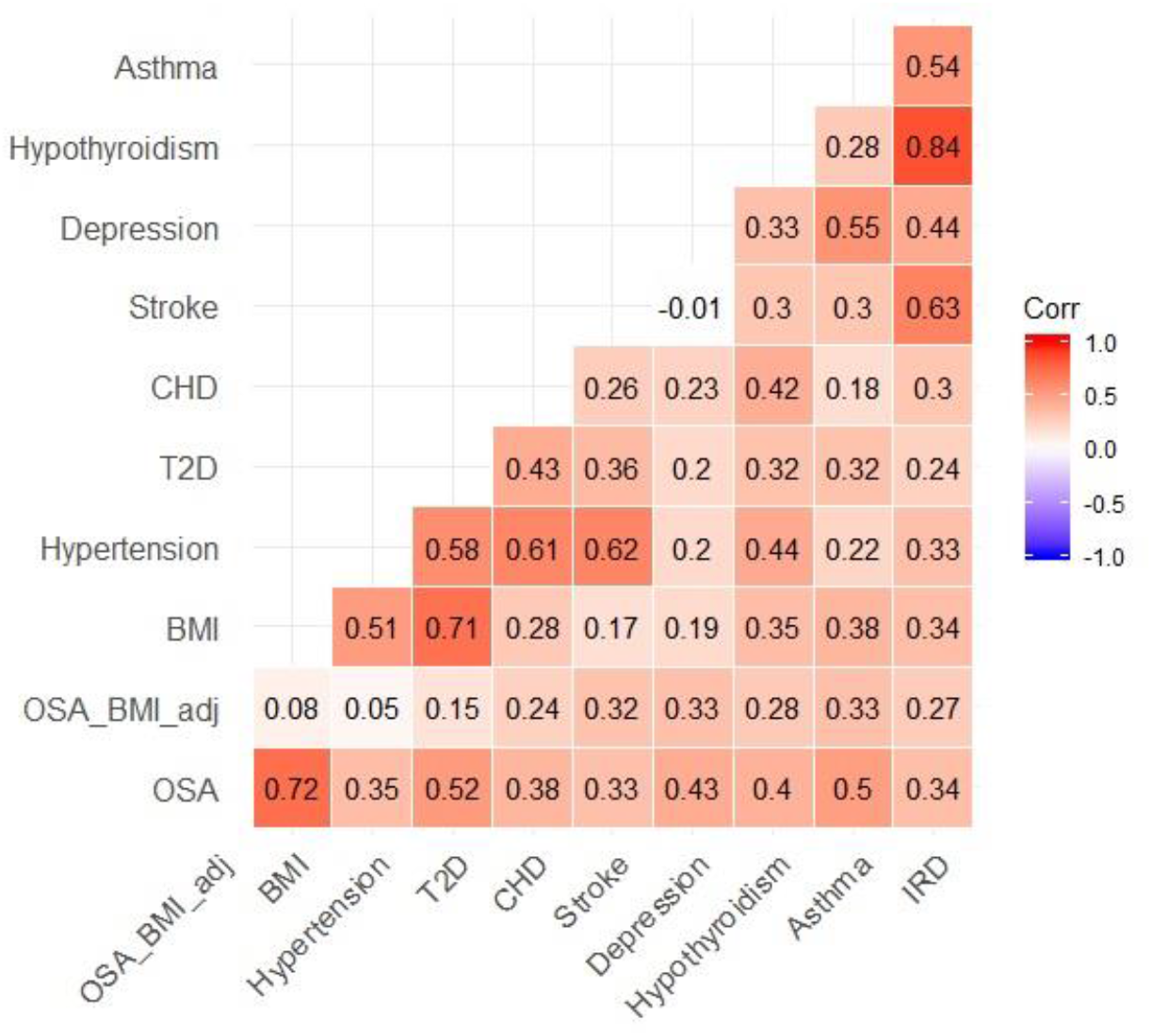
Genetic correlations between obstructive sleep apnoea (OSA), body mass index (BMI) and previously known comorbidities using LD-score regression. Colour-scale represents the strength of the correlation. Correlations between OSA and other traits have been calculated with and without BMI-adjustment. CHD=coronary heart disease, T2D=type 2 diabetes, IRD=inflammatory rheumatic diseases.

To estimate genetic correlations between FinnGen OSA summary statistics and other sleep traits we used UKBB derived summary statistics for sleep variables. We observed genetic correlation with sleep efficiency^13^ rg = −0.31, [−0.44 - −0.17], P=9.80 × 10^−6^) and this was reflected with higher genetic correlation with daytime sleepiness^29^ (rg = 0.44, [0.33-0.54], P=1.27 × 10^−15^). These associations remained significant also after BMI adjustment (rg=-0.19, [−0.36 - −0.03], P= 0.02, rg=0.42, [0.29-0.55], P=1.06 × 10^−10^, respectively). We did not find significant genetic correlations between OSA and sleep duration or chronotype^29^ (**Table 3).**

**Table 3.** Genetic correlations between OSA and other sleep traits

|  | Sleepiness | Sleep duration | Chronotype | Sleep efficiency |
| --- | --- | --- | --- | --- |
| **OSA** | $rg = 0.44 [0.33-0.54]$<br>$p = 1.27 \times 10^{-15}$ | $rg = 0.0096 [-0.085-0.10]$<br>$p = 0.84$ | $rg = -5.0 \times 10^{-4} [-0.079-0.078]$<br>$p = 0.99$ | $rg = -0.31 [-0.44 - -0.17]$<br>$p = 9.80 \times 10^{-6}$ |
| **OSA BMI adjusted** | $rg = 0.42 [0.29-0.55]$<br>$p = 1.06 \times 10^{-10}$ | $rg = 0.078 [-0.031-0.19]$<br>$p = 0.14$ | $rg = -0.063 [-0.154-0.028]$<br>$p = 0.18$ | $rg = -0.19 [-0.36 - -0.03]$<br>$p = 0.02$ |

Summary statistics for sleep traits that were used to calculate the genetic correlations were obtained in previous genome-wide association study (GWAS) from the UK Biobank (UKBB). OSA=obstructive sleep apnoea, BMI=body mass index.

To investigate the biological mechanisms behind OSA, we also examined tissue enrichment of association signals using partitioned heritability analysis using LDSC: an approach which combines data from ENCODE and the GTEx resources^30, 31^ to

FinnGen OSA summary statistics. Concordantly with the association of BMI and cardiometabolic traits, we observed strongest association with cardiovascular tissues and connective and bone tissues (P < 0.05). Furthermore, enrichments with BMI adjusted OSA implicated central nervous system (CNS) as the strongest associating single tissue (P < 0.05) (**Supplementary Figure 3)**.

To test if there is a causal relationship between OSA and its comorbidities, we performed analysis of PRS followed by formal MR analysis using FinnGen OSA summary statistics and independent BMI SNPs^32^. The BMI PRS showed a strong association with OSA risk (**Table 4)** and the individuals in the highest BMI PRS quintile had 1.98-fold increased ([1.88-2.09], P=3.38 × 10^−140^) OSA risk after adjustment for age, sex and 10 first PCs. Similarly, this association was further accentuated in formal MR. We used 64 independent BMI SNPs^32^ as instrumental variables to predict OSA. In line with epidemiological observations and genetic correlation, we discovered a strong causal predictive effect from BMI to OSA (IVW: beta=0.67, P=8.32 × 10^−16^) (**Figure 4, Supplementary Table 3)**.

**Table 4.** BMI’s polygenic risk score predicts OSA

| **BMI_PRS** |  |  |  |
| --- | --- | --- | --- |
|  | OR | CI | p-value |
| BMI_Q1 | - |  | - |
| BMI_Q2 | 1.29 | 1.22-1.36 | $3.49 \times 10^{-19}$ |
| BMI_Q3 | 1.45 | 1.37-1.53 | $5.61 \times 10^{-40}$ |
| BMI_Q4 | 1.61 | 1.53-1.70 | $7.93 \times 10^{-67}$ |
| BMI_Q5 | 1.98 | 1.88-2.09 | $3.38 \times 10^{-140}$ |

Estimated effect coefficients for the body mass index (BMI)'s polygenic risk score as a predictor of OSA. The BMI's polygenic risk score was stratified into quintiles and BMI\_Q5 is the highest quintile. OSA=obstructive sleep apnoea, OR=odds ratio, CI = 95% confidence interval.

**Figure 4.**
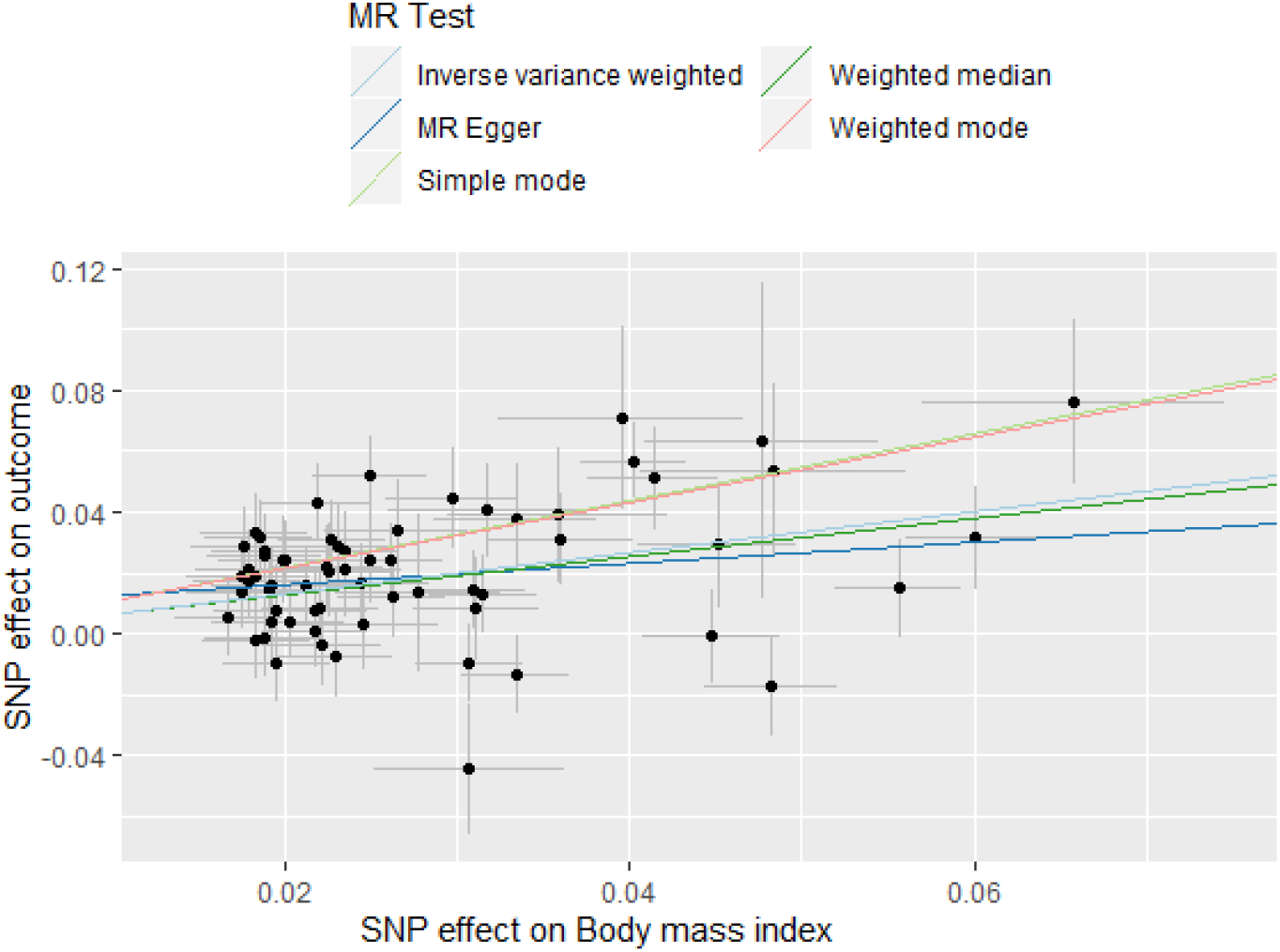
Formal Mendelian randomization (MR) suggesting a strong causal relationship between body mass index (BMI) and obstructive sleep apnoea (OSA) where BMI predicts OSA as an outcome.

### Replication

For each lead variants associated with OSA, we examined the estimates from the additional, comparable cohorts: UKBB, ANDIS and EGCUT. The results were combined using inverse-variance weighted fixed-effect meta-analysis. These additional independent datasets support the role of *FTO* and *GAPVD1* loci in OSA (P < 0.05) (**Supplementary Table 4)**.

## Discussion

In this study, using biobank data of over 217,000 individuals we show that OSA risk has a strong genetic component and identify five genetic loci that are associated with the risk for OSA. Our results show high genetic correlations between OSA and cardiometabolic diseases and risk factors, with strongest connections between OSA and BMI, hypertension, T2D and CHD, which are in line with previous epidemiological and clinical observations. These genetic correlations tracked with phenotypic correlations and comorbidities for OSA. In addition, both our association findings and the MR results support the causal role of obesity in OSA.

These results allow us to draw several conclusions. First, genetic variation plays an important role in development of OSA. This is supported by both the SNP heritability estimates and the associated loci.

Second, our results show that obesity plays a central causal role in the OSA risk. This is supported by high genetic correlations between OSA and BMI. We found that four out of five associated loci were mediated through their associations with BMI. These findings are in line with the finding that weight loss is an important contributor of lowering AHI and the severity of OSA^39, 40^.

Third, we also identified a strong association near *RMST/NEDD1*, which was specific for OSA independent of BMI. The lead SNP associated with antidepressant purchases which may imply that daytime sleepiness caused by OSA together with sleep disturbances may lead to depression and increased antidepressant usage. This is in line with the observation that depression is prevalent among patients with OSA^8^.

Fourth, a strong genetic correlation was observed between OSA and sleep traits, especially with sleepiness and sleep efficiency. These findings highlight the pathological effects of OSA on sleep. As OSA is manageable with Continuous Positive Airway Pressure (CPAP) or oral appliance, these genetic correlations implicate the importance of OSA treatment.

Our study does have some limitations. First, registry-based ascertainment through hospitalisation may miss non-hospitalised cases (false negatives) and treatment information such as CPAP compliance or oral sleep apnoea appliance usage. However, to our knowledge this is the largest number of cases combined for a

GWAS. Second, due to a relatively small number of cases in the replication datasets, our statistical power was limited in the replication analysis. The finding of rs185932673 should be interpreted cautiously as the variant is rare in the Finnish population and the association was not replicated in the other study samples.

Here we present associations between five novel genetic loci and OSA. Our findings highlight the causal links between obesity and OSA but also provide evidence for non-BMI dependent genetic effects. In addition to BMI, we show that genetic effects that modify risk of cardiometabolic diseases, depression, hypothyroidism, asthma and IRD are also correlated with genetic effects for OSA showing that the observed comorbidities between OSA and these diseases may have a joint genetic basis. Our results confirm that OSA is a heterogenic disease with several phenotypes and that implies different approach to OSA management.

## Acknowledgements

We would like to thank all participants of the FinnGen study for their generous participation. We would also like to thank Sari Kivikko for management assistance. Patients and controls in FinnGen provided informed consent for biobank research, based on the Finnish Biobank Act. Alternatively, older research cohorts, collected prior to the start of FinnGen (in August 2017), were collected based on study-specific consents and later transferred to the Finnish biobanks after approval by Valvira, the National Supervisory Authority for Welfare and Health. Recruitment protocols followed the biobank protocols approved by Valvira. The Coordinating Ethics Committee of the Hospital District of Helsinki and Uusimaa (HUS) approved the FinnGen study protocol Nr HUS/990/2017.

The FinnGen study is approved by the Finnish Institute for Health and Welfare (THL), approval numberTHL/2031/6.02.00/2017,amendmentsTHL/1101/5.05.00/2017,THL/341/6.02.00/2018,THL/2222/6.02.00/2018,THL/283/6.02.00/2019,THL/1721/5.05.00/2019,

Digital and population data service agency VRK43431/2017-3, VRK/6909/2018-3, VRK/4415/2019-3 the Social Insurance Institution (KELA) KELA 58/522/2017, KELA 131/522/2018, KELA 70/522/2019, KELA 98/522/2019, and Statistics Finland TK-53-1041-17.

The Biobank Access Decisions for FinnGen samples and data utilised in FinnGen Data Freeze 5 include: THL Biobank BB2017_55, BB2017_111, BB2018_19, BB_2018_34, BB_2018_67, BB2018_71, BB2019_7, BB2019_8, BB2019_26, Finnish Red Cross Blood Service Biobank 7.12.2017, Helsinki Biobank HUS/359/2017, Auria Biobank AB17-5154, Biobank Borealis of Northern Finland_2017_1013, Biobank of Eastern Finland 1186/2018, Finnish Clinical Biobank Tampere MH0004, Central Finland Biobank 1-2017, and Terveystalo Biobank STB 2018001.

This research has been conducted using the UK Biobank Resource under Application Number 22627.

This work was supported by the Academy of Finland Center of Excellence in Complex Disease Genetics [Grant No 312062 to S.R., 312074 to A.P.]; Academy of Finland [Grant No 285380 to S.R, 128650 to A.P, 309643 to H.M.O]; the Finnish Foundation for Cardiovascular Research [to S.R., V.S., and A.P.]; the Sigrid Jusélius Foundation [to S.R. and A.P.]; University of Helsinki HiLIFE Fellow grants 2017-2020 [to S.R.] and Foundation and the Horizon 2020 Research and Innovation Programme [grant number 667301 (COSYN) to A.P.]; Oskar Öfflund foundation and Yrjö Jahnsson foundation [to H.M.O]; The Finnish Dental Society Apollonia [to S.S].

The FinnGen project is funded by two grants from Business Finland (HUS 4685/31/2016 and UH 4386/31/2016) and eleven industry partners (AbbVie Inc, AstraZeneca UK Ltd, Biogen MA Inc, Celgene Corporation, Celgene International II Sàrl, Genentech Inc, Merck Sharp & Dohme Corp, Pfizer Inc., GlaxoSmithKline, Sanofi, Maze Therapeutics Inc., Janssen Biotech Inc). Following biobanks are acknowledged for collecting the FinnGen project samples: Auria Biobank (https://www.auria.fi/biopankki), THL Biobank (https://thl.fi/fi/web/thl-biopankki), Helsinki Biobank (https://www.terveyskyla.fi/helsinginbiopankki), Biobank Borealis of Northern Finland (https://www.oulu.fi/university/node/38474), Finnish Clinical Biobank Tampere (https://www.tays.fi/en-US/Research_and_development/Finnish_Clinical_Biobank_Tampere), Biobank of Eastern Finland (https://ita-suomenbiopankki.fi), Central Finland Biobank (https://www.ksshp.fi/fi-FI/Potilaalle/Biopankki), Finnish Red Cross Blood Service Biobank (https://www.veripalvelu.fi/verenluovutus/biopankkitoiminta) and Terveystalo Biobank (https://www.terveystalo.com/fi/Yritystietoa/Terveystalo-Biopankki/Biopankki/). All Finnish Biobanks are members of BBMRI.fi infrastructure (www.bbmri.fi).

The funders had no role in study design, data collection and analysis, decision to publish, or preparation of the manuscript.

## Author contributions

S.R. and T. P. supervised the study. S.E.R, S.S, H.M.O, M. K. and J.K. performed the statistical and bioinformatics analyses. A.S.H. and T.K. phenotyped study samples. S.S, H.M.O. and S.E.R. wrote the paper with the feedback from all co-authors.

## Data availability

The FinnGen data may be accessed through Finnish Biobanks’ FinnBB portal (www.finbb.fi) and THL Biobank data may be accessed through THL Biobank (https://thl.fi/en/web/thl-biobank).

## Code availability

The full genotyping and imputation protocol for FinnGen is described at https://doi-org.libproxy.helsinki.fi/10.17504/protocols.io.nmndc5e.

## Contributors of FinnGen

### Steering Committee

Aarno Palotie Institute for Molecular Medicine Finland, HiLIFE, University of Helsinki, Finland

Mark Daly Institute for Molecular Medicine Finland, HiLIFE, University of Helsinki, Finland

#### Pharmaceutical companies

Howard Jacob Abbvie, Chicago, IL, United States

Athena Matakidou Astra Zeneca, Cambridge, United Kingdom

Heiko Runz Biogen, Cambridge, MA, United States

Sally John Biogen, Cambridge, MA, United States

Robert Plenge Celgene, Summit, NJ, United States

Mark McCarthy Genentech, San Francisco, CA, United States

Julie Hunkapiller Genentech, San Francisco, CA, United States

Meg Ehm GlaxoSmithKline, Brentford, United Kingdom

Dawn Waterworth GlaxoSmithKline, Brentford, United Kingdom

Caroline Fox Merck, Kenilworth, NJ, United States

Anders Malarstig Pfizer, New York, NY, United States

Kathy Klinger Sanofi, Paris, France

Kathy Call Sanofi, Paris, France

#### University of Helsinki & Biobanks

Tomi Mäkelä HiLIFE, University of Helsinki, Finland, Finland

Jaakko Kaprio Institute for Molecular Medicine Finland, HiLIFE, Helsinki, Finland, Finland

Petri Virolainen Auria Biobank / Univ. of Turku / Hospital District of Southwest Finland, Turku, Finland

Kari Pulkki Auria Biobank / Univ. of Turku / Hospital District of Southwest Finland, Turku, Finland

Terhi Kilpi THL Biobank / Finnish Institute for Health and Welfare Helsinki, Finland

Markus Perola THL Biobank / Finnish Institute for Health and Welfare Helsinki, Finland

Jukka Partanen Finnish Red Cross Blood Service / Finnish Hematology Registry and Clinical Biobank, Helsinki, Finland

Anne Pitkäranta Hospital District of Helsinki and Uusimaa, Helsinki, Finland

Riitta Kaarteenaho Northern Finland Biobank Borealis / University of Oulu / Northern Ostrobothnia Hospital District, Oulu, Finland

Seppo Vainio Northern Finland Biobank Borealis / University of Oulu / Northern Ostrobothnia Hospital District, Oulu, Finland

Kimmo Savinainen Finnish Clinical Biobank Tampere / University of Tampere / Pirkanmaa Hospital District, Tampere, Finland

Veli-Matti Kosma Biobank of Eastern Finland / University of Eastern Finland / Northern Savo Hospital District, Kuopio, Finland

Urho Kujala Central Finland Biobank / University of Jyväskylä / Central Finland Health Care District, Jyväskylä, Finland

#### Other Experts/ Non-Voting Members

Outi Tuovila Business Finland, Helsinki, Finland

Minna Hendolin Business Finland, Helsinki, Finland

Raimo Pakkanen Business Finland, Helsinki, Finland

### Scientific Committee

#### Pharmaceutical companies

Jeff Waring Abbvie, Chicago, IL, United States

Bridget Riley-Gillis Abbvie, Chicago, IL, United States

Athena Matakidou Astra Zeneca, Cambridge, United Kingdom

Heiko Runz Biogen, Cambridge, MA, United States

Jimmy Liu Biogen, Cambridge, MA, United States

Shameek Biswas Celgene, Summit, NJ, United States

Julie Hunkapiller Genentech, San Francisco, CA, United States

Dawn Waterworth GlaxoSmithKline, Brentford, United Kingdom

Meg Ehm GlaxoSmithKline, Brentford, United Kingdom

Dorothee Diogo Merck, Kenilworth, NJ, United States

Caroline Fox Merck, Kenilworth, NJ, United States

Anders Malarstig Pfizer, New York, NY, United States

Catherine Marshall Pfizer, New York, NY, United States

Xinli Hu Pfizer, New York, NY, United States

Kathy Call Sanofi, Paris, France

Kathy Klinger Sanofi, Paris, France

Matthias Gossel Sanofi, Paris, France

#### University of Helsinki & Biobanks

Samuli Ripatti Institute for Molecular Medicine Finland, HiLIFE, University of Helsinki, Helsinki, Finland

Johanna Schleutker Auria Biobank / Univ. of Turku / Hospital District of Southwest Finland, Turku, Finland

Markus Perola THL Biobank / Finnish Institute for Health and Welfare Helsinki, Finland

Mikko Arvas Finnish Red Cross Blood Service / Finnish Hematology Registry and Clinical Biobank, Helsinki, Finland

Olli Carpen Hospital District of Helsinki and Uusimaa, Helsinki, Finland

Reetta Hinttala Northern Finland Biobank Borealis / University of Oulu / Northern Ostrobothnia Hospital District, Oulu, Finland

Johannes Kettunen Northern Finland Biobank Borealis / University of Oulu / Northern Ostrobothnia Hospital District, Oulu, Finland

Reijo Laaksonen Finnish Clinical Biobank Tampere **/** University of Tampere / Pirkanmaa Hospital District, Tampere, Finland

Arto Mannermaa Biobank of Eastern Finland / University of Eastern Finland / Northern Savo Hospital District, Kuopio, Finland

Juha Paloneva Central Finland Biobank / University of Jyväskylä / Central Finland Health Care District, Jyväskylä, Finland

Urho Kujala Central Finland Biobank / University of Jyväskylä / Central Finland Health Care District, Jyväskylä, Finland

#### Other Experts/ Non-Voting Members

Outi Tuovila Business Finland, Helsinki, Finland

Minna Hendolin Business Finland, Helsinki, Finland

Raimo Pakkanen Business Finland, Helsinki, Finland

### Clinical Groups

#### Neurology Group

Hilkka Soininen Northern Savo Hospital District, Kuopio, Finland

Valtteri Julkunen Northern Savo Hospital District, Kuopio, Finland

Anne Remes Northern Ostrobothnia Hospital District, Oulu, Finland

Reetta KälviäinenNorthern Savo Hospital District, Kuopio, Finland

Mikko Hiltunen Northern Savo Hospital District, Kuopio, Finland

Jukka Peltola Pirkanmaa Hospital District, Tampere, Finland

Pentti Tienari Hospital District of Helsinki and Uusimaa, Helsinki, Finland

Juha Rinne Hospital District of Southwest Finland, Turku, Finland

Adam Ziemann Abbvie, Chicago, IL, United States

Jeffrey Waring Abbvie, Chicago, IL, United States

Sahar Esmaeeli Abbvie, Chicago, IL, United States

Nizar Smaoui Abbvie, Chicago, IL, United States

Anne Lehtonen Abbvie, Chicago, IL, United States

Susan Eaton Biogen, Cambridge, MA, United States

Heiko Runz Biogen, Cambridge, MA, United States

Sanni Lahdenperä Biogen, Cambridge, MA, United States

Shameek Biswas Celgene, Summit, NJ, United States

John Michon Genentech, San Francisco, CA, United States

Geoff Kerchner Genentech, San Francisco, CA, United States

Julie Hunkapiller Genentech, San Francisco, CA, United States

Natalie Bowers Genentech, San Francisco, CA, United States

Edmond Teng Genentech, San Francisco, CA, United States

John Eicher Merck, Kenilworth, NJ, United States

Vinay Mehta Merck, Kenilworth, NJ, United States

Padhraig Gormley Merck, Kenilworth, NJ, United States

Kari Linden Pfizer, New York, NY, United States

Christopher Whelan Pfizer, New York, NY, United States

Fanli Xu GlaxoSmithKline, Brentford, United Kingdom

David Pulford GlaxoSmithKline, Brentford, United Kingdom

#### Gastroenterology Group

Martti Färkkilä Hospital District of Helsinki and Uusimaa, Helsinki, Finland

Sampsa Pikkarainen Hospital District of Helsinki and Uusimaa, Helsinki, Finland

Airi Jussila Pirkanmaa Hospital District, Tampere, Finland

Timo Blomster Northern Ostrobothnia Hospital District, Oulu, Finland

Mikko Kiviniemi Northern Savo Hospital District, Kuopio, Finland

Markku Voutilainen Hospital District of Southwest Finland, Turku, Finland Bob Georgantas Abbvie, Chicago, IL, United States

Graham Heap Abbvie, Chicago, IL, United States

Jeffrey Waring Abbvie, Chicago, IL, United States

Nizar Smaoui Abbvie, Chicago, IL, United States

Fedik Rahimov Abbvie, Chicago, IL, United States

Anne Lehtonen Abbvie, Chicago, IL, United States

Keith Usiskin Celgene, Summit, NJ, United States

Joseph Maranville Celgene, Summit, NJ, United States

Tim Lu Genentech, San Francisco, CA, United States

Natalie Bowers Genentech, San Francisco, CA, United States

Danny Oh Genentech, San Francisco, CA, United States

John Michon Genentech, San Francisco, CA, United States

Vinay Mehta Merck, Kenilworth, NJ, United States

Kirsi Kalpala Pfizer, New York, NY, United States

Melissa Miller Pfizer, New York, NY, United States

Xinli Hu Pfizer, New York, NY, United States

Linda McCarthy GlaxoSmithKline, Brentford, United Kingdom

#### Rheumatology Group

Kari Eklund Hospital District of Helsinki and Uusimaa, Helsinki, Finland

Antti Palomäki Hospital District of Southwest Finland, Turku, Finland

Pia Isomäki Pirkanmaa Hospital District, Tampere, Finland

Laura Pirilä Hospital District of Southwest Finland, Turku, Finland

Oili Kaipiainen-Seppänen Northern Savo Hospital District, Kuopio, Finland

Johanna Huhtakangas Northern Ostrobothnia Hospital District, Oulu, Finland

Bob Georgantas Abbvie, Chicago, IL, United States

Jeffrey Waring Abbvie, Chicago, IL, United States

Fedik Rahimov Abbvie, Chicago, IL, United States

Apinya Lertratanakul Abbvie, Chicago, IL, United States

Nizar Smaoui Abbvie, Chicago, IL, United States

Anne Lehtonen Abbvie, Chicago, IL, United States

David Close Astra Zeneca, Cambridge, United Kingdom

Marla Hochfeld Celgene, Summit, NJ, United States

Natalie Bowers Genentech, San Francisco, CA, United States

John Michon Genentech, San Francisco, CA, United States

Dorothee Diogo Merck, Kenilworth, NJ, United States

Vinay Mehta Merck, Kenilworth, NJ, United States

Kirsi Kalpala Pfizer, New York, NY, United States

Nan Bing Pfizer, New York, NY, United States

Xinli Hu Pfizer, New York, NY, United States

Jorge Esparza Gordillo GlaxoSmithKline, Brentford, United Kingdom

Nina Mars Institute for Molecular Medicine Finland, HiLIFE, University of Helsinki, Helsinki, Finland

#### Pulmonology Group

Tarja Laitinen Pirkanmaa Hospital District, Tampere, Finland

Margit Pelkonen Northern Savo Hospital District, Kuopio, Finland

Paula Kauppi Hospital District of Helsinki and Uusimaa, Helsinki, Finland

Hannu Kankaanranta Pirkanmaa Hospital District, Tampere, Finland

Terttu Harju Northern Ostrobothnia Hospital District, Oulu, Finland

Nizar Smaoui Abbvie, Chicago, IL, United States

David Close Astra Zeneca, Cambridge, United Kingdom

Steven GreenbergCelgene, Summit, NJ, United States

Hubert Chen Genentech, San Francisco, CA, United States

Natalie Bowers Genentech, San Francisco, CA, United States

John Michon Genentech, San Francisco, CA, United States

Vinay Mehta Merck, Kenilworth, NJ, United States

Jo Betts GlaxoSmithKline, Brentford, United Kingdom

Soumitra Ghosh GlaxoSmithKline, Brentford, United Kingdom

#### Cardiometabolic Diseases Group

Veikko Salomaa Finnish Institute for Health and Welfare Helsinki, Finland

Teemu Niiranen Finnish Institute for Health and Welfare Helsinki, Finland

Markus Juonala Hospital District of Southwest Finland, Turku, Finland

Kaj Metsärinne Hospital District of Southwest Finland, Turku, Finland

Mika Kähönen Pirkanmaa Hospital District, Tampere, Finland

Juhani Junttila Northern Ostrobothnia Hospital District, Oulu, Finland

Markku Laakso Northern Savo Hospital District, Kuopio, Finland

Jussi Pihlajamäki Northern Savo Hospital District, Kuopio, Finland

Juha Sinisalo Hospital District of Helsinki and Uusimaa, Helsinki, Finland

Marja-Riitta Taskinen Hospital District of Helsinki and Uusimaa, Helsinki, Finland

Tiinamaija Tuomi Hospital District of Helsinki and Uusimaa, Helsinki, Finland

Jari Laukkanen Central Finland Health Care District, Jyväskylä, Finland

Ben Challis Astra Zeneca, Cambridge, United Kingdom

Andrew Peterson Genentech, San Francisco, CA, United States Julie Hunkapiller Genentech, San Francisco, CA, United States

Natalie Bowers Genentech, San Francisco, CA, United States

John Michon Genentech, San Francisco, CA, United States

Dorothee Diogo Merck, Kenilworth, NJ, United States

Audrey Chu Merck, Kenilworth, NJ, United States

Vinay Mehta Merck, Kenilworth, NJ, United States

Jaakko Parkkinen Pfizer, New York, NY, United States

Melissa Miller Pfizer, New York, NY, United States

Anthony Muslin Sanofi, Paris, France

Dawn Waterworth GlaxoSmithKline, Brentford, United Kingdom

#### Oncology Group

Heikki Joensuu Hospital District of Helsinki and Uusimaa, Helsinki, Finland

Tuomo Meretoja Hospital District of Helsinki and Uusimaa, Helsinki, Finland

Olli Carpen Hospital District of Helsinki and Uusimaa, Helsinki, Finland

Lauri Aaltonen Hospital District of Helsinki and Uusimaa, Helsinki, Finland

Annika Auranen Pirkanmaa Hospital District, Tampere, Finland

Peeter Karihtala Northern Ostrobothnia Hospital District, Oulu, Finland

Saila Kauppila Northern Ostrobothnia Hospital District, Oulu, Finland

Päivi Auvinen Northern Savo Hospital District, Kuopio, Finland

Klaus Elenius Hospital District of Southwest Finland, Turku, Finland

Relja Popovic Abbvie, Chicago, IL, United States

Jeffrey Waring Abbvie, Chicago, IL, United States

Bridget Riley-Gillis Abbvie, Chicago, IL, United States

Anne Lehtonen Abbvie, Chicago, IL, United States

Athena Matakidou Astra Zeneca, Cambridge, United Kingdom

Jennifer Schutzman Genentech, San Francisco, CA, United States

Julie Hunkapiller Genentech, San Francisco, CA, United States

Natalie Bowers Genentech, San Francisco, CA, United States

John Michon Genentech, San Francisco, CA, United States

Vinay Mehta Merck, Kenilworth, NJ, United States

Andrey Loboda Merck, Kenilworth, NJ, United States

Aparna Chhibber Merck, Kenilworth, NJ, United States

Heli Lehtonen Pfizer, New York, NY, United States

Stefan McDonough Pfizer, New York, NY, United States

Marika Crohns Sanofi, Paris, France

Diptee Kulkarni GlaxoSmithKline, Brentford, United Kingdom

#### Opthalmology Group

Kai Kaarniranta Northern Savo Hospital District, Kuopio, Finland

Joni Turunen Hospital District of Helsinki and Uusimaa, Helsinki, Finland

Terhi Ollila Hospital District of Helsinki and Uusimaa, Helsinki, Finland

Sanna Seitsonen Hospital District of Helsinki and Uusimaa, Helsinki, Finland

Hannu Uusitalo Pirkanmaa Hospital District, Tampere, Finland

Vesa Aaltonen Hospital District of Southwest Finland, Turku, Finland

Hannele Uusitalo-Järvinen Pirkanmaa Hospital District, Tampere, Finland

Marja LuodonpääNorthern Ostrobothnia Hospital District, Oulu, Finland

Nina Hautala Northern Ostrobothnia Hospital District, Oulu, Finland

Heiko Runz Biogen, Cambridge, MA, United States

Erich Strauss Genentech, San Francisco, CA, United States

Natalie Bowers Genentech, San Francisco, CA, United States

Hao Chen Genentech, San Francisco, CA, United States

John Michon Genentech, San Francisco, CA, United States

Anna Podgornaia Merck, Kenilworth, NJ, United States

Vinay Mehta Merck, Kenilworth, NJ, United States

Dorothee Diogo Merck, Kenilworth, NJ, United States

Joshua Hoffman GlaxoSmithKline, Brentford, United Kingdom

#### Dermatology Group

Kaisa Tasanen Northern Ostrobothnia Hospital District, Oulu, Finland

Laura Huilaja Northern Ostrobothnia Hospital District, Oulu, Finland

Katariina Hannula-Jouppi Hospital District of Helsinki and Uusimaa, Helsinki, Finland

Teea Salmi Pirkanmaa Hospital District, Tampere, Finland

Sirkku Peltonen Hospital District of Southwest Finland, Turku, Finland

Leena Koulu Hospital District of Southwest Finland, Turku, Finland

Ilkka Harvima Northern Savo Hospital District, Kuopio, Finland

Kirsi Kalpala Pfizer, New York, NY, United States

Ying Wu Pfizer, New York, NY, United States

David Choy Genentech, San Francisco, CA, United States

John Michon Genentech, San Francisco, CA, United States

Nizar Smaoui Abbvie, Chicago, IL, United States

Fedik Rahimov Abbvie, Chicago, IL, United States

Anne Lehtonen Abbvie, Chicago, IL, United States

Dawn Waterworth GlaxoSmithKline, Brentford, United Kingdom

### FinnGen Teams

#### Administration Team

Anu Jalanko Institute for Molecular Medicine Finland, HiLIFE, University of Helsinki, Finland

Risto Kajanne Institute for Molecular Medicine Finland, HiLIFE, University of Helsinki, Finland

Ulrike Lyhs Institute for Molecular Medicine Finland, HiLIFE, University of Helsinki, Finland

#### Communication

Mari Kaunisto Institute for Molecular Medicine Finland, HiLIFE, University of Helsinki, Finland

#### Analysis Team

Justin Wade Davis Abbvie, Chicago, IL, United States

Bridget Riley-Gillis Abbvie, Chicago, IL, United States

Danjuma Quarless Abbvie, Chicago, IL, United States

Slavé Petrovski Astra Zeneca, Cambridge, United Kingdom

Jimmy Liu Biogen, Cambridge, MA, United States

Chia-Yen Chen Biogen, Cambridge, MA, United States

Paola Bronson Biogen, Cambridge, MA, United States

Robert Yang Celgene, Summit, NJ, United States

Joseph Maranville Celgene, Summit, NJ, United States

Shameek Biswas Celgene, Summit, NJ, United States

Diana Chang Genentech, San Francisco, CA, United States

Julie Hunkapiller Genentech, San Francisco, CA, United States

Tushar Bhangale Genentech, San Francisco, CA, United States

Natalie Bowers Genentech, San Francisco, CA, United States

Dorothee Diogo Merck, Kenilworth, NJ, United States

Emily Holzinger Merck, Kenilworth, NJ, United States

Padhraig Gormley Merck, Kenilworth, NJ, United States

Xulong Wang Merck, Kenilworth, NJ, United States

Xing Chen Pfizer, New York, NY, United States

Åsa Hedman Pfizer, New York, NY, United States

Kirsi Auro GlaxoSmithKline, Brentford, United Kingdom

Clarence Wang Sanofi, Paris, France

Ethan Xu Sanofi, Paris, France

Franck Auge Sanofi, Paris, France

Clement Chatelain Sanofi, Paris, France

Mitja Kurki Institute for Molecular Medicine Finland, HiLIFE, University of Helsinki, Finland /

Broad Institute, Cambridge, MA, United States

Samuli Ripatti Institute for Molecular Medicine Finland, HiLIFE, University of Helsinki, Finland

Mark Daly Institute for Molecular Medicine Finland, HiLIFE, University of Helsinki, Finland

Juha Karjalainen Institute for Molecular Medicine Finland, HiLIFE, University of Helsinki, Finland /

Broad Institute, Cambridge, MA, United States

Aki Havulinna Institute for Molecular Medicine Finland, HiLIFE, University of Helsinki, Finland

Anu Jalanko Institute for Molecular Medicine Finland, HiLIFE, University of Helsinki, Finland

Kimmo Palin University of Helsinki, Helsinki, Finland

Priit Palta Institute for Molecular Medicine Finland, HiLIFE, University of Helsinki, Finland

Pietro Della Briotta Parolo Institute for Molecular Medicine Finland, HiLIFE, University of Helsinki, Finland Wei Zhou Broad Institute, Cambridge, MA, United States

Susanna Lemmelä Institute for Molecular Medicine Finland, HiLIFE, University of Helsinki, Finland

Manuel Rivas University of Stanford, Stanford, CA, United States

Jarmo Harju Institute for Molecular Medicine Finland, HiLIFE, University of Helsinki, Finland

Aarno Palotie Institute for Molecular Medicine Finland, HiLIFE, University of Helsinki, Finland

Arto Lehisto Institute for Molecular Medicine Finland, HiLIFE, University of Helsinki, Finland

Andrea Ganna Institute for Molecular Medicine Finland, HiLIFE, University of Helsinki, Finland

Vincent Llorens Institute for Molecular Medicine Finland, HiLIFE, University of Helsinki, Finland

Antti Karlsson Auria Biobank / Univ. of Turku / Hospital District of Southwest Finland, Turku, Finland

Kati Kristiansson THL Biobank / Finnish Institute for Health and Welfare Helsinki, Finland

Mikko Arvas Finnish Red Cross Blood Service / Finnish Hematology Registry and Clinical Biobank, Helsinki, Finland

Kati Hyvärinen Finnish Red Cross Blood Service / Finnish Hematology Registry and Clinical Biobank, Helsinki, Finland

Jarmo Ritari Finnish Red Cross Blood Service / Finnish Hematology Registry and Clinical Biobank, Helsinki, Finland

Tiina Wahlfors Finnish Red Cross Blood Service / Finnish Hematology Registry and Clinical Biobank, Helsinki, Finland

Miika Koskinen Hospital District of Helsinki and Uusimaa, Helsinki, Finland BB/HUS/Univ Hosp Districts

Olli Carpen Hospital District of Helsinki and Uusimaa, Helsinki, Finland BB/HUS/Univ Hosp Districts

Johannes Kettunen Northern Finland Biobank Borealis / University of Oulu / Northern Ostrobothnia Hospital District, Oulu, Finland

Katri Pylkäs Northern Finland Biobank Borealis / University of Oulu / Northern Ostrobothnia Hospital District, Oulu, Finland

Marita Kalaoja Northern Finland Biobank Borealis / University of Oulu / Northern Ostrobothnia

Hospital District, Oulu, Finland

Minna Karjalainen Northern Finland Biobank Borealis / University of Oulu / Northern Ostrobothnia Hospital District, Oulu, Finland

Tuomo Mantere Northern Finland Biobank Borealis / University of Oulu / Northern Ostrobothnia

Hospital District, Oulu, Finland

Eeva Kangasniemi Finnish Clinical Biobank Tampere **/** University of Tampere / Pirkanmaa Hospital District, Tampere, Finland

Sami Heikkinen Biobank of Eastern Finland / University of Eastern Finland / Northern Savo Hospital District, Kuopio, Finland

Arto Mannermaa Biobank of Eastern Finland / University of Eastern Finland / Northern Savo Hospital District, Kuopio, Finland

Eija Laakkonen Central Finland Biobank / University of Jyväskylä / Central Finland Health Care District, Jyväskylä, Finland

Juha Kononen Central Finland Biobank / University of Jyväskylä / Central Finland Health Care District, Jyväskylä, Finland

#### Sample Collection Coordination

Anu Loukola Hospital District of Helsinki and Uusimaa, Helsinki, Finland

#### Sample Logistics

Päivi Laiho THL Biobank / Finnish Institute for Health and Welfare Helsinki, Finland

Tuuli Sistonen THL Biobank / Finnish Institute for Health and Welfare Helsinki, Finland

Essi Kaiharju THL Biobank / Finnish Institute for Health and Welfare Helsinki, Finland

Markku Laukkanen THL Biobank / Finnish Institute for Health and Welfare Helsinki, Finland

Elina Järvensivu THL Biobank / Finnish Institute for Health and Welfare Helsinki, Finland

Sini Lähteenmäki THL Biobank / Finnish Institute for Health and Welfare Helsinki, Finland

Lotta Männikkö THL Biobank / Finnish Institute for Health and Welfare Helsinki, Finland

Regis Wong THL Biobank / Finnish Institute for Health and Welfare Helsinki, Finland

#### Registry Data Operations

Kati Kristiansson THL Biobank / Finnish Institute for Health and Welfare Helsinki, Finland

Hannele Mattsson THL Biobank / Finnish Institute for Health and Welfare Helsinki, Finland

Susanna Lemmelä Institute for Molecular Medicine Finland, HiLIFE, University of Helsinki, Finland

Tero Hiekkalinna THL Biobank / Finnish Institute for Health and Welfare Helsinki, Finland

Manuel González Jiménez. THL Biobank / Finnish Institute for Health and Welfare Helsinki, Finland

#### Genotyping

Kati Donner Institute for Molecular Medicine Finland, HiLIFE, University of Helsinki, Finland

#### Sequencing Informatics

Priit Palta Institute for Molecular Medicine Finland, HiLIFE, University of Helsinki, Finland

Kalle Pärn Institute for Molecular Medicine Finland, HiLIFE, University of Helsinki, Finland

Javier Nunez-Fontarnau Institute for Molecular Medicine Finland, HiLIFE, University of Helsinki, Finland

#### Data Management and IT Infrastructure

Jarmo Harju Institute for Molecular Medicine Finland, HiLIFE, University of Helsinki, Finland

Elina Kilpeläinen Institute for Molecular Medicine Finland, HiLIFE, University of Helsinki, Finland

Timo P. Sipilä Institute for Molecular Medicine Finland, HiLIFE, University of Helsinki, Finland

Georg Brein Institute for Molecular Medicine Finland, HiLIFE, University of Helsinki, Finland

Alexander Dada Institute for Molecular Medicine Finland, HiLIFE, University of Helsinki, Finland

Ghazal Awaisa Institute for Molecular Medicine Finland, HiLIFE, University of Helsinki, Finland

Anastasia Shcherban Institute for Molecular Medicine Finland, HiLIFE, University of Helsinki, Finland

Tuomas Sipilä Institute for Molecular Medicine Finland, HiLIFE, University of Helsinki, Finland

#### Clinical Endpoint Development

Hannele Laivuori Institute for Molecular Medicine Finland, HiLIFE, University of Helsinki, Finland

Aki Havulinna Institute for Molecular Medicine Finland, HiLIFE, University of Helsinki, Finland

Susanna Lemmelä Institute for Molecular Medicine Finland, HiLIFE, University of Helsinki, Finland

Tuomo Kiiskinen Institute for Molecular Medicine Finland, HiLIFE, University of Helsinki, Finland

#### Trajectory Team

Tarja Laitinen Pirkanmaa Hospital District, Tampere, Finland

Harri Siirtola University of Tampere, Tampere, Finland

Javier Gracia Tabuenca University of Tampere, Tampere, Finland

#### Biobank Directors

Lila Kallio Auria Biobank, Turku, Finland

Sirpa Soini THL Biobank, Helsinki, Finland

Jukka Partanen Blood Service Biobank, Helsinki, Finland

Kimmo Pitkänen Helsinki Biobank, Helsinki, Finland

Seppo Vainio Northern Finland Biobank Borealis, Oulu, Finland

Kimmo Savinainen Tampere Biobank, Tampere, Finland

Veli-Matti Kosma Biobank of Eastern Finland, Kuopio, Finland

Teijo Kuopio Central Finland Biobank, Jyväskylä, Finlan

## References

1. Senaratna, C. V. et al. Prevalence of obstructive sleep apnea in the general population: A systematic review. Sleep Med. Rev. 34, 70–81 (2017).

2. Young, T., Evans, L., Finn, L. & Palta, M. Estimation of the clinically diagnosed proportion of sleep apnea syndrome in middle-aged men and women. Sleep 20, 705–706 (1997).

3. Finkel, K. J. et al. Prevalence of undiagnosed obstructive sleep apnea among adult surgical patients in an academic medical center. Sleep Med. 10, 753–758 (2009).

4. Kohler, M. & Stradling, J. R. Mechanisms of vascular damage in obstructive sleep apnea. Nat. Rev. Cardiol. 7, 677–685 (2010).

5. Young, T., Skatrud, J. & Peppard, P. E. Risk factors for obstructive sleep apnea in adults. JAMA 291, 2013–2016 (2004).

6. Strausz, S. et al. Obstructive sleep apnoea and the risk for coronary heart disease and type 2 diabetes: a longitudinal population-based study in Finland. BMJ Open 8, e022752–2018 (2018).

7. Fu, Y. et al. Meta-analysis of all-cause and cardiovascular mortality in obstructive sleep apnea with or without continuous positive airway pressure treatment. Sleep Breath 21, 181–189 (2017).

8. BaHammam, A. S. et al. Comorbid depression in obstructive sleep apnea: an under-recognized association. Sleep Breath 20, 447–456 (2016).

9. Bahammam, S. A., Sharif, M. M., Jammah, A. A. & Bahammam, A. S. Prevalence of thyroid disease in patients with obstructive sleep apnea. Respir. Med. 105, 1755–1760 (2011).

10. Kong, D. L. et al. Association of Obstructive Sleep Apnea with Asthma: A Meta-Analysis. Sci. Rep. 7, 4088–017 (2017).

11. Taylor-Gjevre, R. M., Nair, B. V. & Gjevre, J. A. Obstructive sleep apnoea in relation to rheumatic disease. Rheumatology (Oxford) 52, 15–21 (2013).

12. Redlund-Johnell, I. Upper airway obstruction in patients with rheumatoid arthritis and temporomandibular joint destruction. Scand. J. Rheumatol. 17, 273–279 (1988).

13. Farias Tempaku, P. et al. Genome-wide association study reveals two novel risk alleles for incident obstructive sleep apnea in the EPISONO cohort. Sleep Med. 66, 24–32 (2019).

14. Cade, B. E. et al. Genetic Associations with Obstructive Sleep Apnea Traits in Hispanic/Latino Americans. Am. J. Respir. Crit. Care Med. 194, 886–897 (2016).

15. Chen, H. et al. Multiethnic Meta-Analysis Identifies RAI1 as a Possible Obstructive Sleep Apnea-related Quantitative Trait Locus in Men. Am. J. Respir. Cell Mol. Biol. 58, 391–401 (2018).

16. Campos, A. I. et al. Insights into the aetiology of snoring from observational and genetic investigations in the UK Biobank. Nat. Commun. 11, 817–020 (2020).

17. Veatch, O. J. et al. Characterization of genetic and phenotypic heterogeneity of obstructive sleep apnea using electronic health records. BMC Med. Genomics 13, 105–020 (2020).

18. Tolonen, H. et al. The validation of the Finnish Hospital Discharge Register and Causes of Death Register data on stroke diagnoses. Eur. J. Cardiovasc. Prev. Rehabil. 14, 380–385 (2007).

19. Romero-Corral, A., Caples, S. M., Lopez-Jimenez, F. & Somers, V. K. Interactions between obesity and obstructive sleep apnea: implications for treatment. Chest 137, 711–719 (2010).

20. Konecny, T., Kara, T. & Somers, V. K. Obstructive sleep apnea and hypertension: an update. Hypertension 63, 203–209 (2014).

21. Wang, X., Bi, Y., Zhang, Q. & Pan, F. Obstructive sleep apnoea and the risk of type 2 diabetes: a meta-analysis of prospective cohort studies. Respirology 18, 140–146 (2013).

22. Dong, J. Y., Zhang, Y. H. & Qin, L. Q. Obstructive sleep apnea and cardiovascular risk: meta-analysis of prospective cohort studies. Atherosclerosis 229, 489–495 (2013).

23. Zhou, W. et al. Efficiently controlling for case-control imbalance and sample relatedness in large-scale genetic association studies. Nat. Genet. 50, 1335–1341 (2018).

24. Bulik-Sullivan, B. K. et al. LD Score regression distinguishes confounding from polygenicity in genome-wide association studies. Nat. Genet. 47, 291–295 (2015).

25. 1000 Genomes Project Consortium et al. An integrated map of genetic variation from 1,092 human genomes. Nature 491, 56–65 (2012).

26. International HapMap 3 Consortium et al. Integrating common and rare genetic variation in diverse human populations. Nature 467, 52–58 (2010).

27. Lane, J. M. et al. Genome-wide association analyses of sleep disturbance traits identify new loci and highlight shared genetics with neuropsychiatric and metabolic traits. Nat. Genet. 49, 274–281 (2017).

28. Jones, S. E. et al. Genome-Wide Association Analyses in 128,266 Individuals Identifies New Morningness and Sleep Duration Loci. PLoS Genet. 12, e1006125 (2016).

29. Jones, S. E. et al. Genetic studies of accelerometer-based sleep measures yield new insights into human sleep behaviour. Nat. Commun. 10, 1585–019 (2019).

30. Finucane, H. K. et al. Partitioning heritability by functional annotation using genome-wide association summary statistics. Nat. Genet. 47, 1228–1235 (2015).

31. Finucane, H. K. et al. Heritability enrichment of specifically expressed genes identifies disease-relevant tissues and cell types. Nat. Genet. 50, 621–629 (2018).

32. Locke, A. E. et al. Genetic studies of body mass index yield new insights for obesity biology. Nature 518, 197–206 (2015).

33. Ge, T., Chen, C. Y., Ni, Y., Feng, Y. A. & Smoller, J. W. Polygenic prediction via Bayesian regression and continuous shrinkage priors. Nat. Commun. 10, 1776–019 (2019).

34. Paternoster, L., Tilling, K. & Davey Smith, G. Genetic epidemiology and Mendelian randomization for informing disease therapeutics: Conceptual and methodological challenges. PLoS Genet. 13, e1006944 (2017).

35. Watanabe, K., Taskesen, E., van Bochoven, A. & Posthuma, D. Functional mapping and annotation of genetic associations with FUMA. Nat. Commun. 8, 1826–017 (2017).

36. Pulit, S. L. et al. Meta-analysis of genome-wide association studies for body fat distribution in 694 649 individuals of European ancestry. Hum. Mol. Genet. 28, 166–174 (2019).

37. Hoffmann, T. J. et al. A Large Multiethnic Genome-Wide Association Study of Adult Body Mass Index Identifies Novel Loci. Genetics 210, 499–515 (2018).

38. Frayling, T. M. et al. A common variant in the FTO gene is associated with body mass index and predisposes to childhood and adult obesity. Science 316, 889–894 (2007).

39. Joosten, S. A. et al. Improvement in Obstructive Sleep Apnea With Weight Loss is Dependent on Body Position During Sleep. Sleep 40, 10.1093/sleep/zsx047 (2017).

40. Myllymaa, K. et al. Effect of oxygen desaturation threshold on determination of OSA severity during weight loss. Sleep Breath 20, 33–42 (2016).

