## Supplementary material for "Genetic analysis of obstructive sleep apnoea discovers a strong association with cardiometabolic health"

**Supplementary information**

FinnGen samples were genotyped with Illumina and Affymetrix arrays (Thermo Fisher Scientific, Santa Clara, CA, USA). Genotype calls were made with GenCall and zCall algorithms for Illumina and AxiomGT1 algorithm for Affymetrix chip genotyping data. Genotyping data produced with previous chip platforms were lifted over to build version 38 (GRCh38/hg38) following the protocol described here: dx.doi.org/10.17504/protocols.io.nqtddwn. Samples with sex discrepancies, high genotype missingness (> 5%), excess heterozygosity (+-4SD) and non-Finnish ancestry were removed. Variants with high missingness (> 2%), deviation from Hardy–Weinberg equilibrium ($P$ < 1e-6) and low minor allele count (MAC < 3) were removed. Pre-phasing of genotyped data was performed with Eagle 2.3.5 (https://data.broadinstitute.org/alkesgroup/Eagle/) with the default parameters, except the number of conditioning haplotypes was set to 20,000. Imputation was carried out by using the population-specific Sequencing Initiative Suomi (SISu) v3 imputation reference panel with Beagle 4.1 (version 08Jun17.d8b, https://faculty.washington.edu/browning/beagle/b4_1.html) as described in the following protocol: [dx.doi.org/10.17504/protocols.io.nmndc5e]. SISu v3 imputation reference panel was developed using the high-coverage (25–30x) whole-genome sequencing data generated at the Broad Institute of MIT and Harvard and at the McDonnell Genome Institute at Washington University, USA; and jointly processed at the Broad Institute. Variant callset was produced with Genomic Analysis Toolkit (GATK) HaplotypeCaller algorithm by following GATK best-practices for variant calling. Genotype-, sample- and variant-wise quality control was applied in an iterative manner by using the Hail framework v0.1 (https://Github.com/hail-is/hail/releases/tag/0.2.13, http://Doi.org/10.5281/zenodo.2646680). The resulting high-quality whole genome sequencing data for 3775 individuals were phased with Eagle 2.3.5 as described above. Post-imputation quality control involved excluding variants with INFO score < 0.7.

### Supplementary Table 1. The main findings of the previous GWAS studies

| 1st author | Trait | Sample size | Original GWAS finding | | Corresponding finding in FinnGen | Corresponding finding in FinnGen (BMI adjusted) |
|---|---|---|---|---|---|---|
| Tempaku F[13] | Obstructive sleep apnoea trait (apnoea hypopnea index, change over time) | 706 | rs12415421 | beta=0.28<br>p=3.4 x $10^{-8}$ | beta=0.032<br>p=0.38 | beta=0.048<br>p=0.26 |
| | | | rs4731117 | beta=0.28<br>p=4.4 x $10^{-8}$ | beta=0.014<br>p=0.37 | beta= 0.019<br>p=0.29 |
| Chen H[14] | Obstructive sleep apnoea trait (apnoea hypopnea index) NREM AHI in men | Total: 19,744<br>Men: 6,737 | rs12936587 | beta=0.12<br>p=1.7 × $10^{-8}$ | beta=0.0023<br>p=0.86 | beta=0.0097<br>p=0.53 |
| Cade B[15] | Obstructive sleep apnoea trait (apnea hypopnea index, average respiratory event duration) | 12,558 | rs116791765 | beta=-0.32<br>p=1. 9 x $10^{-8}$ | Not defined in the FinnGen data | Not defined in the FinnGen data |
| | | | rs35424364 | beta=0.03<br>p=4.9 x $10^{-8}$ | beta=-0.014<br>p=0.51 | beta=-0.0034<br>p=0.89 |

The main results of the previous genome-wide association studies (GWAS) and comparison to the FinnGen data findings. BMI=body mass index.

### Supplementary Table 2. ICD-codes for OSA and comorbidities

| Phenotype endpoint | ICD-10 | ICD-9 | ICD-8 |
|---|---|---|---|
| **OSA** | G47.3 | 3472 | |
| **HYPERTENSION** | I10-I13, I15, I67.4 | 4019X, 4029A, 4029B, 4039A, 4040A, 4059A, 4059B, 4372A, 4059X | 400, 401, 402, 403, 404 |
| **T2D** | E11 | 250A | |
| **CHD** | I20.0, I21, I22 | 410, 4110 | 410, 4110 |
| **STROKE** | I61, I63, I64 | 431, 4330A, 4331A, 4339A, 4340A, 4341A, 4349A, 436 | 431, 433, 434, 436 |
| **DEPRESSION** | F32, F33 | 2961, 2968 | 790,20, 298,0 |
| **HYPOTHYROIDISM** | E00, E01, E02, E03.0-E03.5, E03.8, E03.9 | 243, 2443, 2448, 2449, 2448A, 2448B | 243, 244 |
| **ASTHMA** | J45, J46 | 493 | 493 |
| **IRD** | M05, J99.0, M06.0, M30-M35, M45, M08.0, L40.5 | 7140A, 7140B, 7141, 7100, 7431, 7101, 7340, 7200, 7143A, 6960A | 712,10, 712,4, 712,0, 696,00 |

By combining codes from different registries, we generate phenotype “endpoints”. Finnish national version for each ICD-codes were used. These ICD-code criteria are all so-called regular expressions. OSA=obstructive sleep apnoea, T2D=type 2 diabetes, CHD=coronary heart disease, IRD= inflammatory rheumatic diseases.

**Supplementary Table 3. Mendelian randomization suggesting a strong causal relationship between BMI and OSA.**

| Method | number of SNPs | beta | Se | p-value |
|---|---|---|---|---|
| MR Egger | 64 | 0.35 | 0.24 | 0.15 |
| Weighted median | 64 | 0.64 | 0.11 | $1.53 \times 10^{-8}$ |
| Inverse variance weighted | 64 | 0.67 | 0.08 | $8.32 \times 10^{-16}$ |
| Simple mode | 64 | 1.09 | 0.30 | $6.42 \times 10^{-4}$ |
| Weighted mode | 64 | 1.08 | 0.26 | $1.25 \times 10^{-4}$ |

Mendelian randomization (MR) analysis uses 64 independent BMI associated SNPs[32] as an instrumental variable to predict OSA.

**Supplementary Table 4. Replication of the lead variants**

| **RSID** | **G47.3 OSA UKBB** | **G47.3 OSA ANDIS** | **G47.3 OSA EGCUT** | **G47.3 OSA Combined** |
|---|---|---|---|---|
| **case/control** | 4471/403723 | 947/9829 | 4930/61056 | 10348/474608 |
| **rs9937053** | OR=1.12 [1.07-1.17]<br>P=$5.5 \times 10^{-7}$ | OR=1.13 [1.03-1.24]<br>P=0.01 | OR=1.06 [1.02-1.11]<br>P=$6.55 \times 10^{-3}$ | OR=1.09 [1.06-1.12]<br>P=$2.68 \times 10^{-9}$ |
| **rs10507084** | OR=1.07 [0.98-1.17]<br>P=0.15 | OR=0.89 [0.73-1.06]<br>P=0.18 | OR=1.01 [0.94-1.09]<br>P=0.80 | OR=1.02 [0.96-1.08]<br>P=0.51 |
| **rs185932673** | OR=0.96 [0.73-1.26]<br>P=0.74 | Not defined in ANDIS | OR=1.09[0.84-1.43]<br>P=0.52 | OR=1.02[0.84-1.23]<br>P=0.82 |
| **rs4837016** | OR=0.97 [0.93-1.01]<br>P=0.16 | OR=0.87 [0.79-0.95]<br>P=$4.6 \times 10^{-3}$ | OR=0.98 [0.94-1.02]<br>P=0.32 | OR=0.96 [0.94-0.99]<br>P=0.01 |
| **rs10928560** | OR=1.00 [0.94-1.06]<br>P=0.94 | OR=1.01[0.90-1.15]<br>P=0.38 | OR=1.01 [0.96-1.07]<br>P=0.60 | OR=1.01 [0.97-1.05]<br>P= 0.57 |

Inverse-variance weighted meta-analysis combining the results of the replication cohorts of the main FinnGen findings considering obstructive sleep apnoea (OSA). OR=odds ratio, [95% confidence interval], UKBB = UK Biobank, ANDIS = All New Diabetics in Scania, EGCUT = Estonian Genome Center - University of Tartu.

### Supplementary Figure 1.

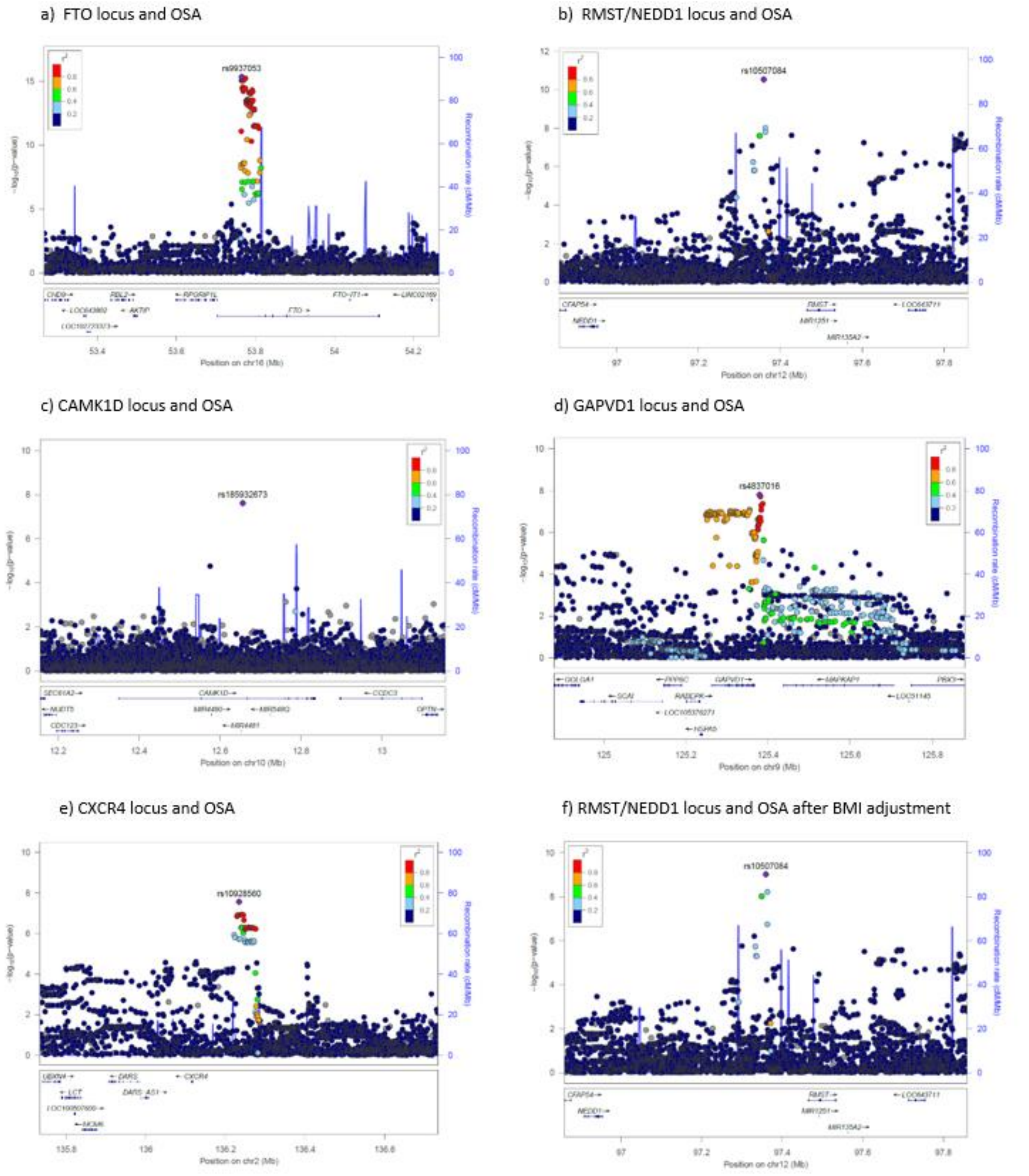


Regional plots of 5 confirmed associations. Locus Zoom plots a-f show associated P-values on the $-\log_{10}$ scale on the vertical axis, and the chromosomal position along the horizontal axis. Purple diamonds indicate SNP at each locus with the strongest associated evidence. LD ($r^2$ values) between the lead SNP and the other SNPs are indicated by colour. *FTO*=Fat mass and obesity-associated protein, *RMST*=Rhabdomyosarcoma 2 associated transcript / *NEDD1*=NEDD1 gamma-tubulin ring complex targeting factor, *CAMK1D*=Calcium/calmodulin-dependent protein kinase ID, *GAPVD1*= GTPase activating protein and VPS9 domains 1, *CXCR4*=C-X-C Motif chemokine receptor 4.

**Supplementary Figure 2.**

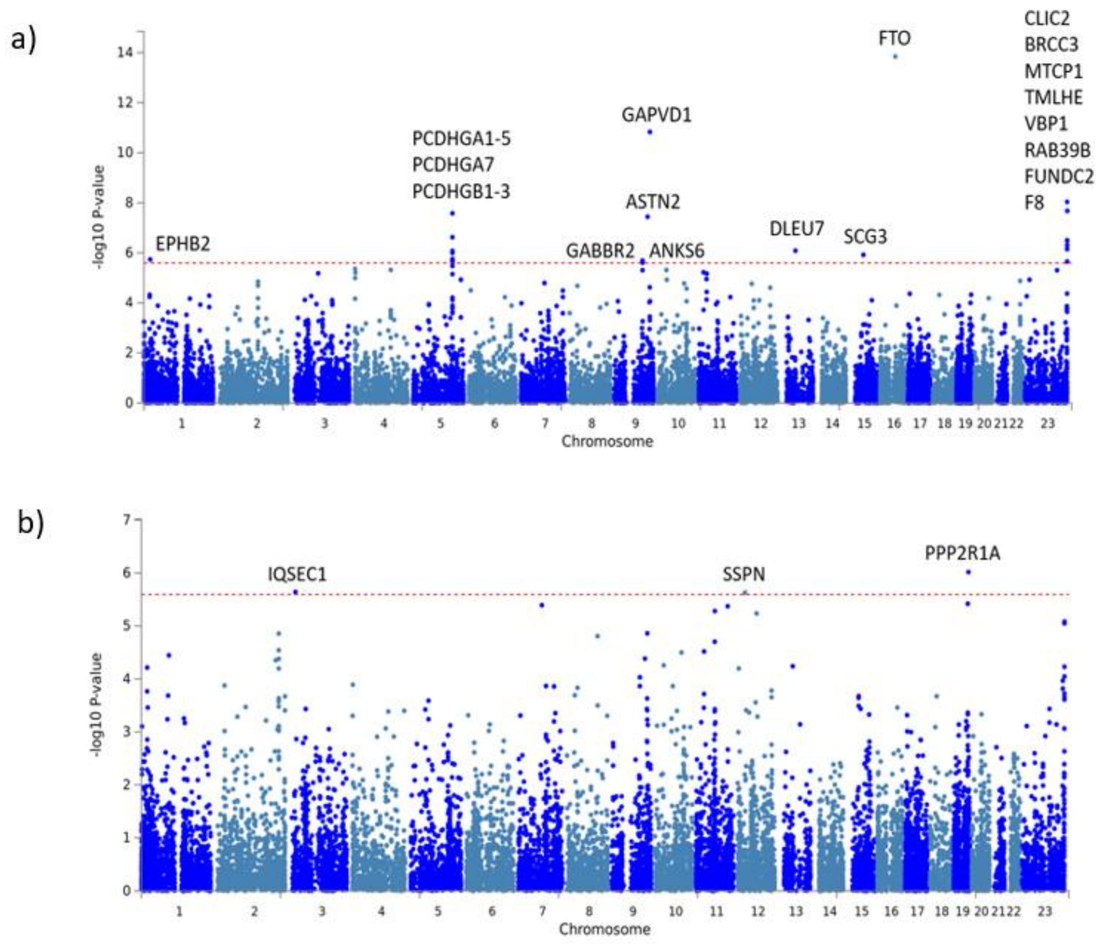


a) Manhattan plot of the gene-based test as computed by MAGMA. Single nucleotide polymorphisms (SNP)s were mapped to 19,651 protein coding genes. Significance Bonferroni corrected threshold was defined at P = 0.05/19,651 = 2.54 $\times 10^{-6}$. Primarily the same genes were identified as in single variant associations. For each annotated gene x-axis shows the chromosomal position while y-axis shows the −log10(P) value. *EPHB2*=Ephrin type-B receptor 2, *PCDHGA*=Protocadherin gamma subfamily A, *PCDHGB*=Protocadherin gamma subfamily B, *GAPVD1*=GTPase activating protein and VPS9 domains 1, *ASTN2*= Astrotactin 2, *GABBR2*=Gamma-aminobutyric acid type A receptor subunit rho2, *ANKS6*=Ankyrin repeat and sterile alpha motif domain containing 6, *DLEU7*=Deleted in lymphocytic leukemia 7, *SCG3*=Secretogranin III, *FTO*=Fat mass and obesity-associated protein, *CLIC2*=Chloride intracellular channel 2, *BRCC3*=BRCA1/BRCA2-containing complex subunit 3, *MTCP1*=Mature T cell proliferation 1, *TMLHE*=Trimethyllysine hydroxylase, epsilon, *VBP1*= VHL binding protein 1, *RAB39B*=RAB39B, member RAS oncogene family, *FUNDC2*=FUN14 domain containing 2 and *F8*= Coagulation factor VIII.

b) Manhattan plot of the gene-based test as computed by MAGMA using body mass index (BMI) adjusted GWAS data. Single nucleotide polymorphisms (SNP)s were mapped to 19,651 protein coding genes. Significance Bonferroni corrected threshold was defined at P = 0.05/19,651 = 2.54 $\times 10^{-6}$. For each annotated gene x-axis shows the chromosomal position while y-axis shows the −log10(P) value. *IQSEC1*= IQ motif and sec7 domain arfGEF 1, *SSPN*=Sarcospan, *PPP2R1A*= Protein phosphatase 2 scaffold subunit aalpha.

### Supplementary Figure 3.

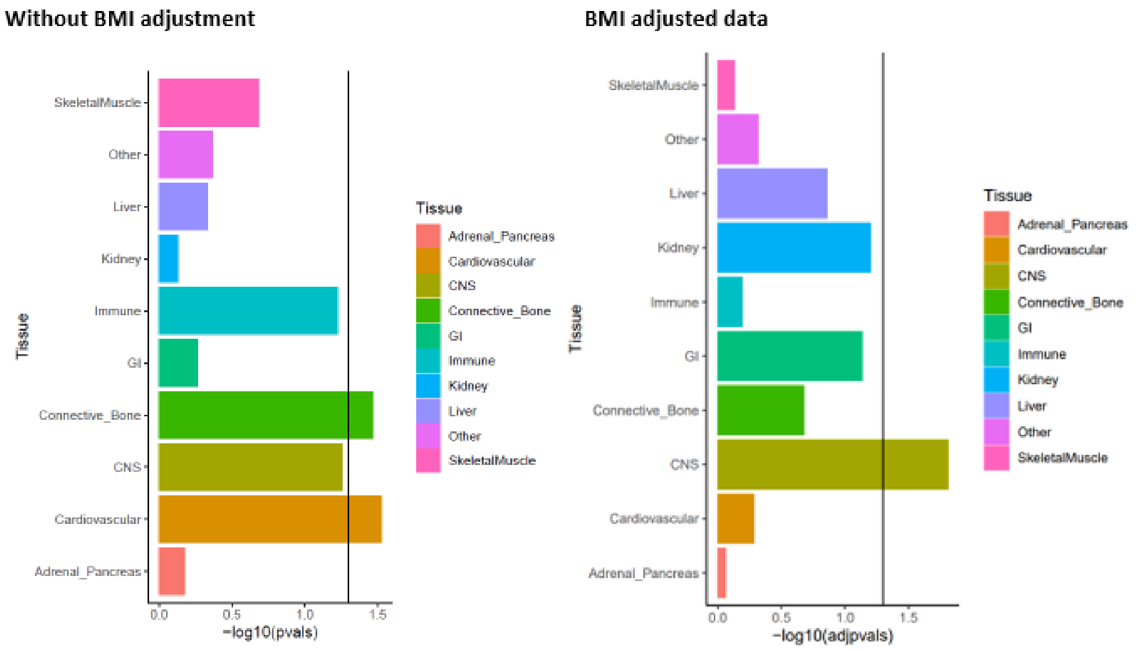


Tissue specific enrichment analysis. Stratified LD score regression based on 1000 Genomes Project phase 1. LD was calculated by each tissue types. Each bar represents -log10 p-value for enrichment and computed for obstructive sleep apnoea (OSA) and body mass index (BMI) adjusted OSA.  CNS=central nervous system, GI=gastrointestinal.
